# LFMD: detecting low-frequency mutations in high-depth genome sequencing data without molecular tags

**DOI:** 10.1101/617381

**Authors:** Rui Ye, Xuehan Zhuang, Jie Ruan, Yanwei Qi, Yitai An, Jiaming Xu, Timothy Mak, Xiao Liu, Xiuqing Zhang, Huanming Yang, Xun Xu, Larry Baum, Chao Nie, Pak Chung Sham

## Abstract

As next-generation sequencing (NGS) and liquid biopsy become more prevalent in research and in the clinic, there is an increasing need for better methods to reduce cost and improve sensitivity and specificity of low-frequency mutation detection (where the Alternative Allele Frequency, or AAF, is less than 1%). Here we propose a likelihood-based approach, called Low-Frequency Mutation Detector (LFMD), which combines the advantages of duplex sequencing (DS) and the bottleneck sequencing system (BotSeqS) to maximize the utilization of duplicate reads. Compared with the existing state-of-the-art methods, DS, Du Novo, UMI-tools, and Unified Consensus Maker, our method achieves higher sensitivity, higher specificity (< 4 × 10^−10^ errors per base sequenced) and lower cost (reduced by ~70% at best) without involving additional experimental steps, customized adapters or molecular tags. LFMD is useful in areas where high precision is required, such as drug resistance prediction and cancer screening. As an example of LFMD’s applications, mitochondrial heterogeneity analysis of 28 human brain samples across different stages of Alzheimer’s Disease (AD) showed that the canonical oxidative damage related mutations, C:G>A:T, are significantly increased in the mid-stage group. This is consistent with the Mitochondrial Free Radical Theory of Aging, suggesting that AD may be linked to the aging of brain cells induced by oxidative damage.

## Background

Low-frequency mutations (LFMs) are defined as somatic mutations with allele frequency lower than 1% in an individual’s DNA or mixed DNA. LFMs can indicate early stages of diseases (such as cancer[1–3], Alzheimer’s Disease (AD)[4] and Parkinson’s Disease (PD)[5]), study aging[5, 6], identify disease-causing variants[7], predict potential drug resistance[8], diagnose mitochondrial disease before tri-parental *in vitro* fertilization[9], and track the mutational spectrum of viral genomes, malignant lesions, and somatic tissues[8, 10]. To effectively improve signal-to-noise ratio (SNR) and detect LFMs, researchers have developed methods with stringent thresholds, complex experimental procedures[4, 11], single cell sequencing[12–15], circle sequencing[16], o2n-seq[17], Sef-SeqS[18, 19], and more precise analytic models[6, 20–22]. The bottleneck sequencing system[23] (BotSeqS) and duplex sequencing[24, 25] (DS) utilize duplicate reads generated by polymerase chain reaction (PCR), which are discarded by other methods, to achieve much higher accuracy. However, current methods still have some limitations in detecting LFMs.

### Disadvantages of single cell sequencing

For single cell sequencing, DNA extraction is laborious and exacting, with point mutations and copy number biases introduced during the amplification of small amounts of fragile DNA. To increase specificity, only variants shared by at least two cells are accepted as true variants[15]. At present, this method is not cost-efficient and cannot be used in large-scale clinical applications because a large number of single cells need to be sequenced to identify rare mutations.

### Disadvantages of circle sequencing and o2n-seq

Circle sequencing and o2n-seq only utilize single-strand DNA templates, so their specificities are limited by the error rate of PCR. Circle sequencing controls errors to a rate as low as 7.6 × 10^−6^ per base sequenced[16] and o2n-seq’s error rate ranges from 10^−5^ to 10^−8^ per base sequenced[26], while DS can achieve 4 × 10^−10^ errors per base sequenced[24].

### Disadvantages of BotSeqS

In contrast, BotSeqS uses endogenous molecular tags to group reads from the same DNA template and construct double-strand consensus reads. As a result, it can detect very rare mutations (<10^−6^) while being cheap enough to sequence the whole human genome[23]. However, it requires highly diluted DNA templates before PCR amplification to reduce endogenous tag conflicts and ensure sufficient sequencing of each DNA template. Thus, it has a high specificity with poor sensitivity. Also, it discards clonal variants and small insertions/deletions (indels) to limit false positives.

### Disadvantages of DS

DS ligates exogenous random molecular tags (also known as barcode, unique molecular identifier, UID or UMI) to both ends of each DNA template prior to PCR amplification. Although sensitive and accurate, much sequencing data is wasted on sequencing tags, fixed sequences and a large proportion of read families that contain less than three read pairs. The errors on molecular tags lead to false construction of read families, smaller read families, and lower utilization rate of sequencing data. In addition, since tags are synthesized with customized adapters, batch effects may occur during DNA library construction.

### Disadvantages of tag clustering

To solve the problem induced by the errors on tags, multiple methods (e.g. UMI-tools) have been developed to cluster similar tags[27–29], where one or two mismatches are allowed in merging two tags. Although tag clustering does improve the sensitivity, it is still not a straightforward way to solve the problem. Inappropriate tag clustering might occur because unstable synthesis of random tags can result in distinct but similar tags.

### A new approach

To avoid the aforementioned problems, we present here a new, efficient approach that combines the advantages of BotSeqS and DS. The method uses a likelihood-based model[6, 20, 30] to dramatically reduce endogenous tag conflicts. Then it groups reads into read families and constructs double-strand consensus reads to detect ultra-rare mutations accurately while maximizing the utilization of duplicate read pairs. This simplifies the DNA sequencing procedure, saves data and cost, achieves higher sensitivity and specificity, and can be used in whole genome sequencing if coupled with a dilution step prior to PCR amplification. In addition, our new method offers a statistical solution to the problem of PCR duplication in the basic analysis pipeline of NGS data and can improve sensitivity and specificity of other variant calling algorithms without specific experimental designs. As the price of sequencing is falling, the depth and the rate of PCR duplication are rising. The method we present here might help deal with such high-depth data more accurately and efficiently.

Since the LFMs of multiple samples need to be compared collectively to unveil the relationship between LFMs and phenotypes, the detection sensitivities in all samples should be estimated carefully, based on the differences in sequencing depth, initial DNA concentration, mutation frequency, PCR amplification efficiency, double-strand consensus sequence (DCS) depth, etc. Meanwhile, it is known that these factors significantly affect the sensitivity of LFM detection. If the results are compared directly without adjustment, systematic bias may conceal the real differences between samples. Therefore, it is essential to estimate the sensitivity of the method for each position at which any mutation was detected.

## Results

### Overview of LFMD

The strategy of LFMD (pronounced “leaf mid”) is quite intuitive, like “looking for a needle in a haystack”: split the haystack into smaller ones, and then look for the needle in the smaller haystacks. In LFMD, the needle is a DNA fragment that has a mutation and several PCR duplicates; the clustering of read pairs with the same genome coordinates is splitting the haystack into smaller ones. Because the allele frequency in the smaller haystack is much higher than that in the original haystack, it is much easier to find the needle; i.e., sensitivity is increased.

To group reads from the same DNA template, the simplest idea is to group properly mapped reads with the same coordinates (i.e., chromosome, start position, and end position) because random shearing of DNA molecules can provide natural differences, called endogenous tags, between templates. A group of reads is called a read family. However, as the length of each DNA template is only approximately determined, random shearing cannot provide enough differences to distinguish each DNA template. Thus, it is common that two original DNA templates share the same coordinates. If two or more DNA templates shared the same coordinates, and their reads are grouped into a single read family, it is difficult to determine, using only their frequencies as a guide, whether an allele is a potential error or a mutation. Thus, BotSeqS introduced a strategy of dilution before PCR amplification to dramatically reduce the number of DNA templates in order to reduce the probability of endogenous tag conflicts. And DS introduced exogenous molecular tags before PCR amplification to dramatically increase the differences between templates. Thus, BotSeqS sacrifices sensitivity and DS sequences extra data: the tags.

Here we introduce a third strategy to eliminate tag conflicts. It is a likelihood-based approach based on an intuitive hypothesis: that if reads of two or more DNA templates group together due to endogenous tags, a true allele’s frequency in this read family is high enough to distinguish the allele from background sequencing errors. The overview of the LFMD pipeline is shown in Figure 1.

**Figure 1.**
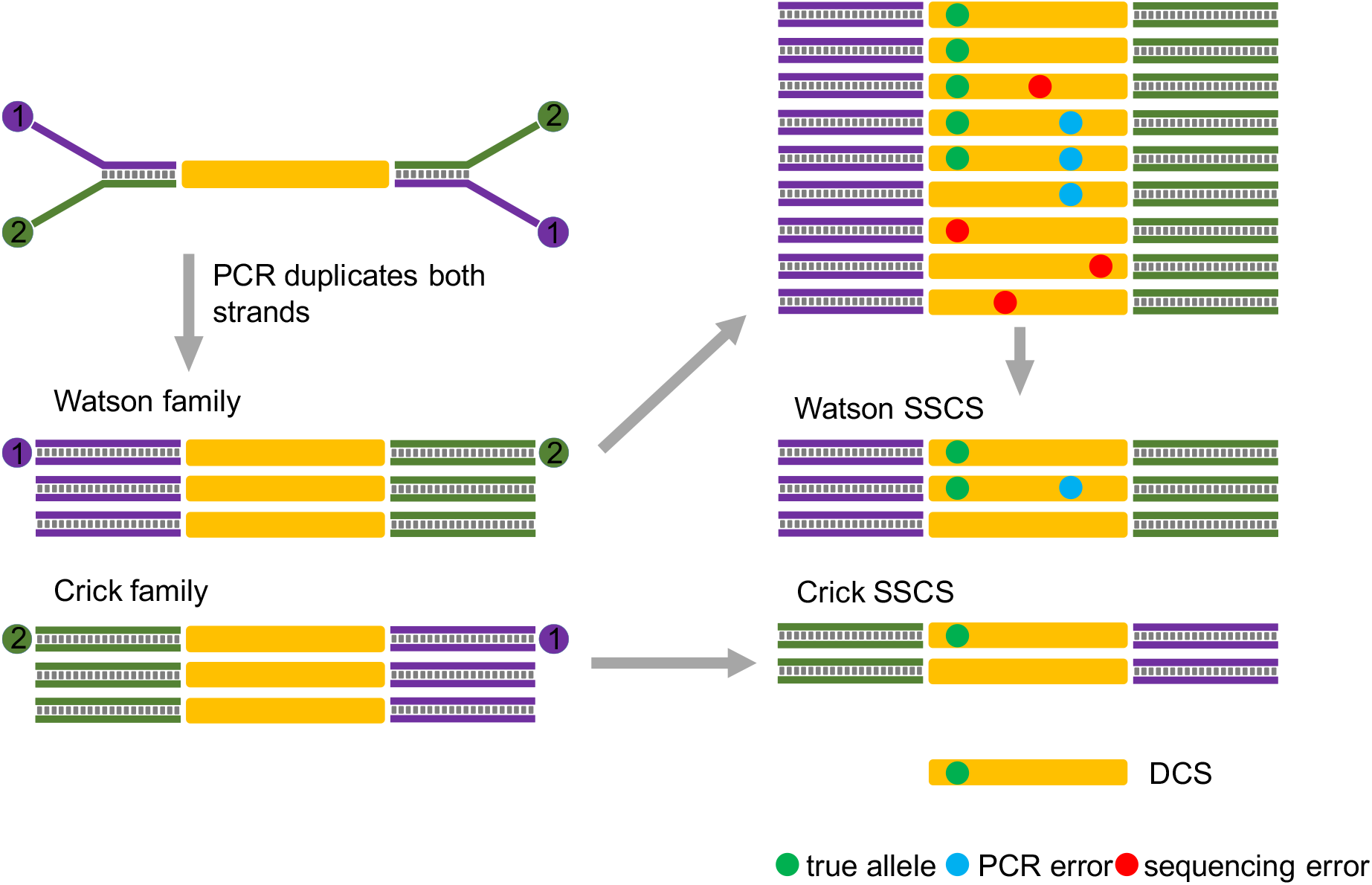
Overview of LFMD. The Y-shaped adapters determine read 1 (purple bar) and 2 (green bar). The directions of reads determine +/− strands. So, after the first cycle of the PCR amplification, the Watson and Crick families are well defined. Then within a read family, true alleles (green dots) and accumulated PCR errors (blue dots) are detected via the likelihood-base model and given a combined error rate. Sequencing errors (red dots) are eliminated. Combining single-strand consensus sequences (SSCSs) of paired read families, high-quality double-strand consensus sequences (DCSs) with estimated error rates are generated and used in the downstream analysis.

### Performance of the method on simulated data

We modified Python scripts developed by the Du Novo[31] team to simulate mixed double-strand sequencing data. A total of 800 single nucleotide variants (SNVs), 120 insertions (INSs), and 120 deletions (DELs) were simulated in the human mitochondrial genome. Although the simulation may not be entirely realistic, the results are still useful to evaluate the performance and the potential drawbacks of LFMD, Du Novo[31], DS, UMI-tools, and Unified Consensus Maker (UniC, an advanced version of DS). Because the underlying true mutations are known, the true positives and false positives can be calculated (Figures 2 and 3). Among these tools, DS and UniC are from the same lab. UniC is an updated version of DS. DS uses the genome positions to groups read pairs based on their genome positions and considers molecular tags later, while UniC groups read pairs into read families based on molecular tags prior to read alignment. UMI-tools is used to cluster molecular tags to increase the size of read families. Du Novo uses the same strategy as UniC but requires an extremely large amount of memory, and its output consensus reads have embedded false indels which cause many false positives, several magnitudes higher than that of other tools. So, in the following comparison, we do not compare Du Novo with other tools. With different sequencing depths—33k, 65k, and 130k— and distinct mutation frequencies, LFMD consistently has the best performance in terms of sensitivity and specificity (Figures 2 and 3).

**Figure 2.**
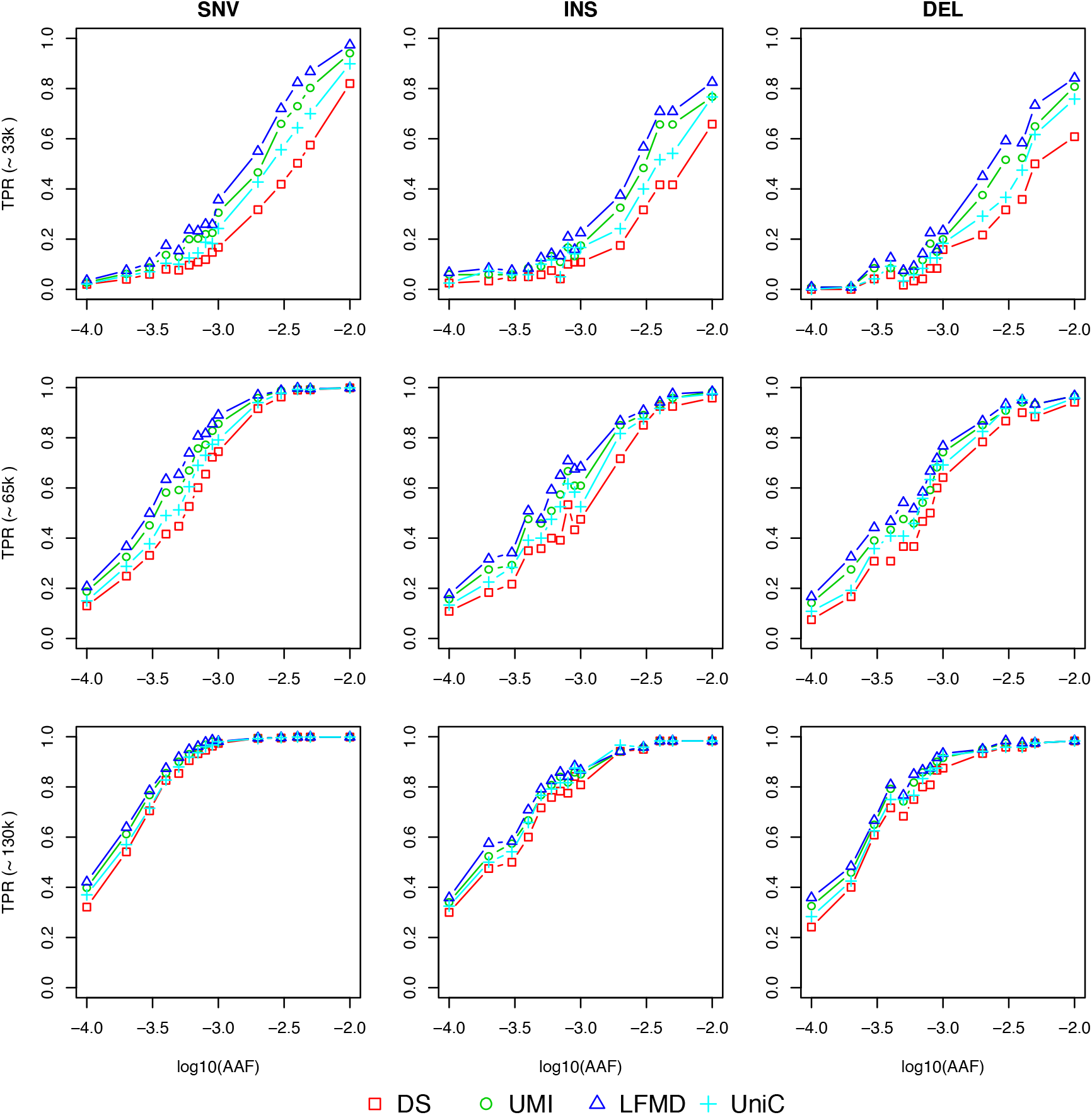
True positive rates of four tools on simulated data with depth around 33k, 65k, and 130k. DS, duplex sequencing; UMI, UMI-tools; UniC, Unified Consensus Maker; SNV, single nucleotide variant; INS, small insertion; DEL, small deletion; TPR, true positive rate; AAF, alternative allele frequency;

**Figure 3.**
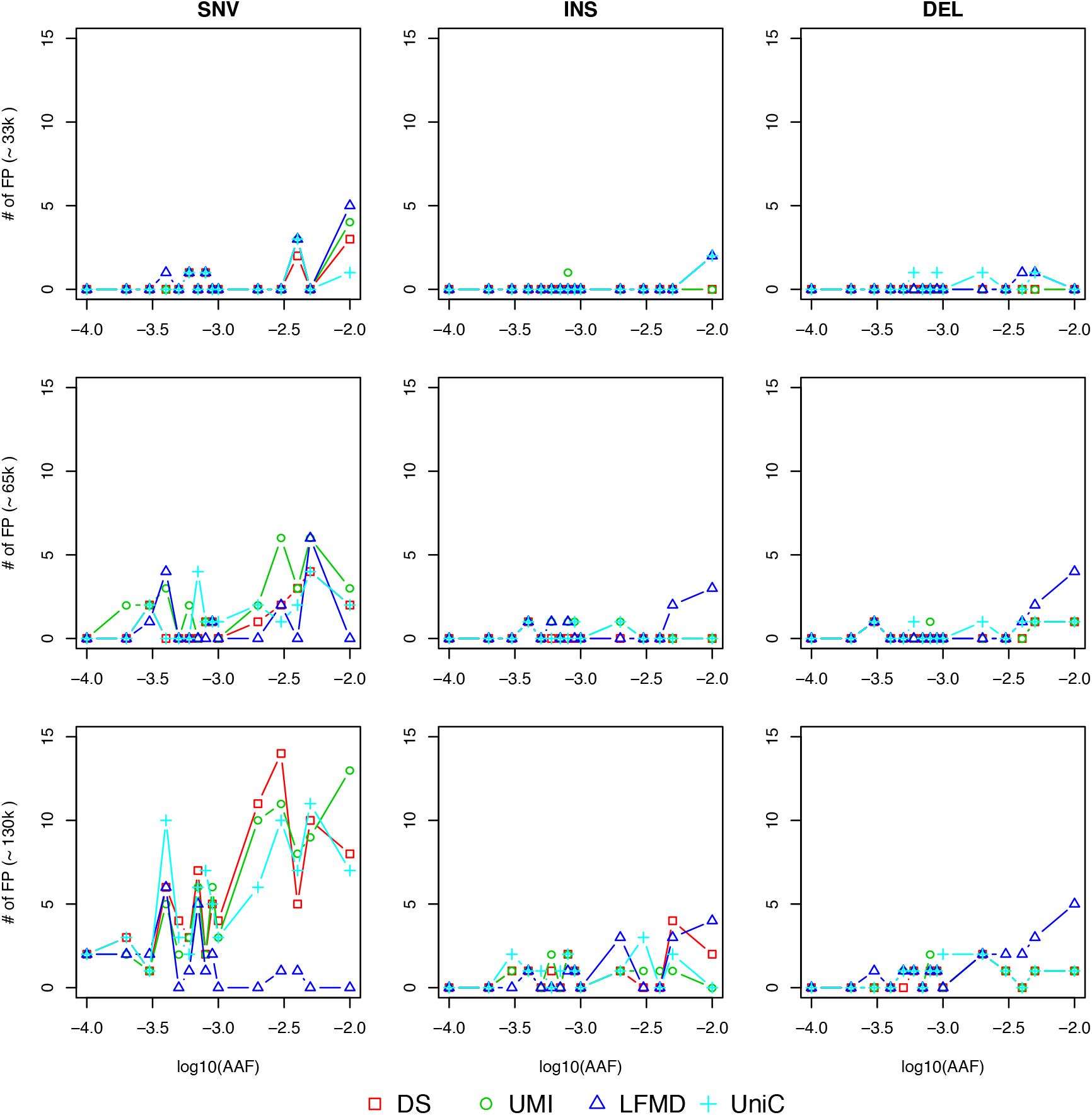
The number of false positives of four tools on simulated data with depth around 33k, 65k, and 130k. DS, duplex sequencing; UMI, UMI-tools; UniC, Unified Consensus Maker; SNV, single nucleotide variant; INS, small insertion; DEL, small deletion; FP, false positive; AAF, alternative allele frequency.

### Performance of the method on real data

To evaluate the performance of LFMD on real data, we used consistent parameters to rerun the whole pipelines of DS, UMI-tools (UMI), Unified Consensus Maker (UniC), and LFMD in a mouse mitochondrial DNA (mtDNA) dataset: SRR1613972 (Figure 4a). The pipelines are designed to only allow fundamental differences between the four strategies and fix other parameters in the analysis pipeline. Thus, the comparison between the results indicates the underlying differences of the tools. To reduce the influence of mapping quality, we only use unique, properly mapped, read pairs to call mutations.

**Figure 4.**
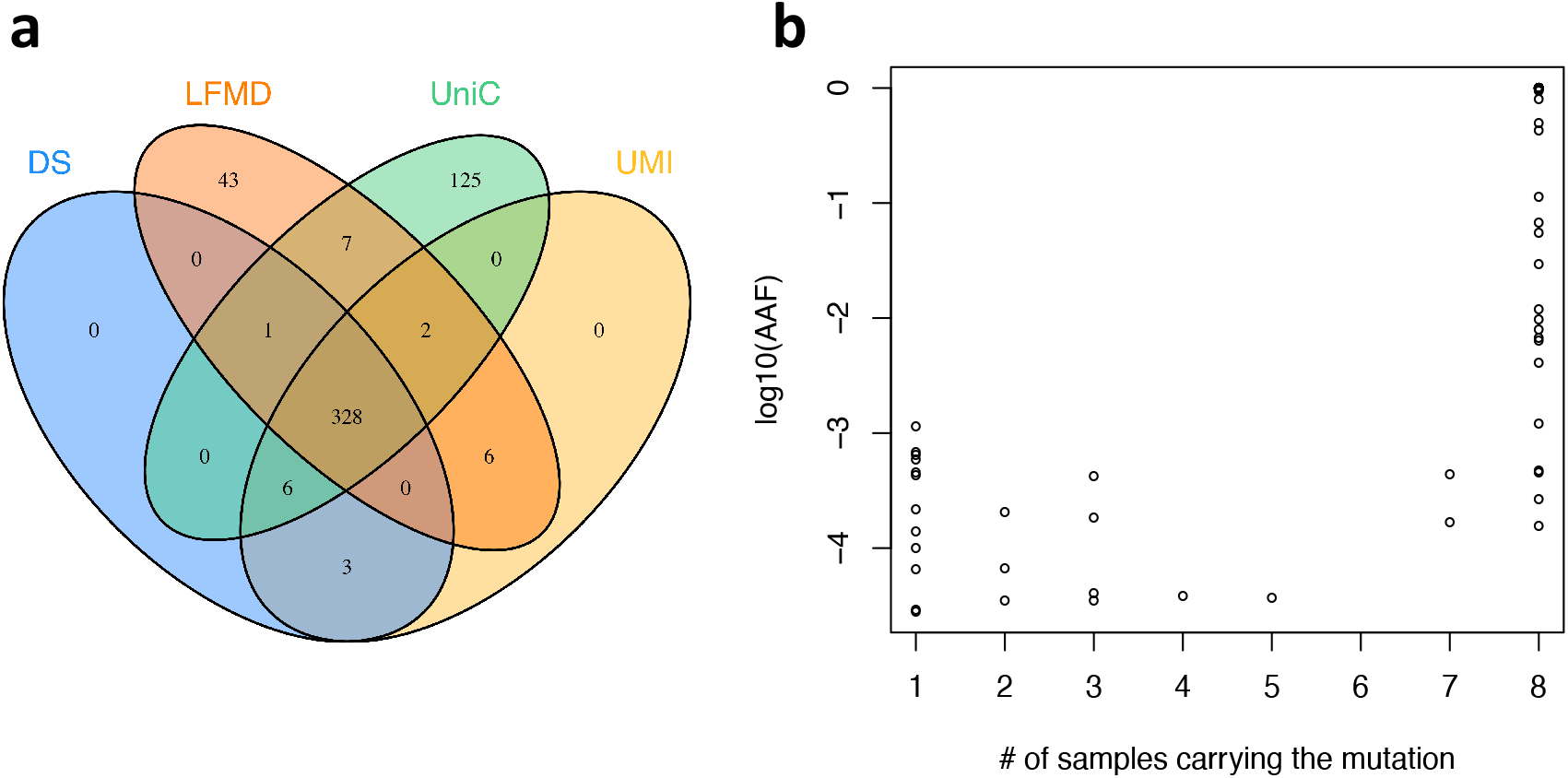
Venn diagram and robustness of LFMD on real data. (a) Venn diagram of mutations detected by four tools in mouse mtDNA. (b) Distribution of mutations found in mtDNA of the Yan Huang cell line. DS, duplex sequencing; UMI, UMI-tools; UniC, Unified Consensus Maker; AAF, alternative allele frequency.

We investigated the discordant mutations one by one. Three mutations only detected by DS and UMI-tools may be false positives due to poor mapping qualities (MT:7G>A and MT:3418T>del) and low sequencing qualities (MT:5462T>G). The mapping errors are from the following: 1) DS and UMI-tools assigned the highest base quality for all bases of DCSs[24] even if the base was N and 2) DS and UMI-tools introduced some Ns in DCS reads because the minimum proportion of “true” bases in a read family is set to 0.7 by default (it can be set to 0.5 to improve sensitivity). Forty-three mutations only detected by LFMD were due to four technical reasons: 1) sequencing and PCR errors on tags led to smaller read families in DS, decreasing its sensitivity; 2) some low complexity tags were discarded; 3) the tools except LFMD assumed “true” mutations should occupy most reads (proportion >= 0.7) in the read family, although this assumption is unreasonable considering that PCR errors occur in the first PCR cycle; 4) potential false positives of LFMD occurred near indels due to mapping errors.

LFMD displayed a type of error which led to 6 possible false negatives compared to other tools. This error may occur when a short indel error in a read family splits the big read family into smaller ones, reducing the ability to detect mutations in the read family. UniC detected some mutations that were not detected by other tools. This may be related to the fact that UniC tended to report more false positives in the simulated data due to barcode conflict.

### Consistency evaluation using 8 technical replicates of Yan Huang (YH) cell line

We sequenced the YH cell line (passage 19) in 8 independent experiments to evaluate the stability of LFMD (see Materials for details). The results are highly consistent in terms of numbers of mutations detected (range 61~68), shown in Figure 4b and Table 1. Under the hypothesis that true mutations should be identified from at least two samples, we detected 68 “true” mutations, with the mean true positive rate (TPR) and false discovery rate (FDR) around 91.4% and 2.4%, respectively. Due to the high specificity and low sensitivity of LFMD in simulated data, it is very unlikely that any of the mutations are false positives, and the above TPR and FDR only indicate the consistency of parallel experiments.

**Table 1.** The number of mutations found in mtDNA of 8 YH cell lines. Under the hypothesis that true mutations should be identified from at least two samples, we detected 68 “true” mutations and then calculated TP, FP, TPR, and FDR.

| Samples | # of mutations | TP | FP | TPR | FDR |
| --- | --- | --- | --- | --- | --- |
| L01_501 | 64 | 63 | 1 | 92.7% | 1.56% |
| L01_502 | 68 | 67 | 1 | 98.5% | 1.47% |
| L01_503 | 62 | 62 | 0 | 91.2% | 0.00% |
| L01_504 | 65 | 63 | 2 | 92.7% | 3.08% |
| L01_505 | 62 | 60 | 2 | 88.2% | 3.23% |
| L01_506 | 61 | 59 | 2 | 86.8% | 3.28% |
| L01_507 | 65 | 62 | 3 | 91.2% | 4.62% |
| L01_508 | 62 | 61 | 1 | 89.7% | 1.61% |
| Mean | 63.6 | 62.1 | 1.50 | 91.4% | 2.4% |
| SD | 2.3 | 2.4 | 0.93 | 3.6% | 1.5% |

TP, true positive; FP, false positive; TPR, true positive rate; FDR, false discovery rate; SD, standard deviation.

### Mitochondrial DNA Data from 28 Samples with Alzheimer’s Disease

The dataset used in this section was provided by Prof. Scott R. Kennedy[4]. They studied point mutations and indels in mtDNA of 28 samples with AD and reported the possibility that the accumulation of low frequency mutations in mtDNA plays a key role in the occurrence and development of AD. They also claimed that mtDNA mutations are inconsistent with oxidative damage. The samples are flash frozen tissues obtained from the hippocampus and parietal lobes of human brain. According to the criteria from DSM-IV and Braak Staging[32], cases were stratified into three groups: ND-Low, ND-High, and D-High, representing early-, mid-, and late-stages, respectively. This dataset was reanalysed with four tools: DS, UMI-tools (UMI), Unified Consensus Maker (UniC), and LFMD. Due to the higher sensitivity, LFMD detected more mutations than other tools, especially for C:G>A:T mutations in the ND-High group (Supplementary Figure S7).

After adjustment for sensitivity, the canonical signature of oxidative damage on DNA[33–35], C:G>A:T, was significantly more common (p<0.01) in the ND-High group than in the ND-Low group (Figure 5). This is inconsistent with the main conclusion of Hoekstra et al.’s paper[4] (the origin of the dataset), but consistent with other papers which focus on high frequency mutations in mtDNA[36–38]. In conclusion, a more sensitive tool, coupled with appropriate adjustments, can help uncover the true relationship between LFMs and phenotypes. The low frequency C:G>A:T mutations in mtDNA are markers of oxidative DNA damage and play a role in AD pathogenesis.

**Figure 5.**
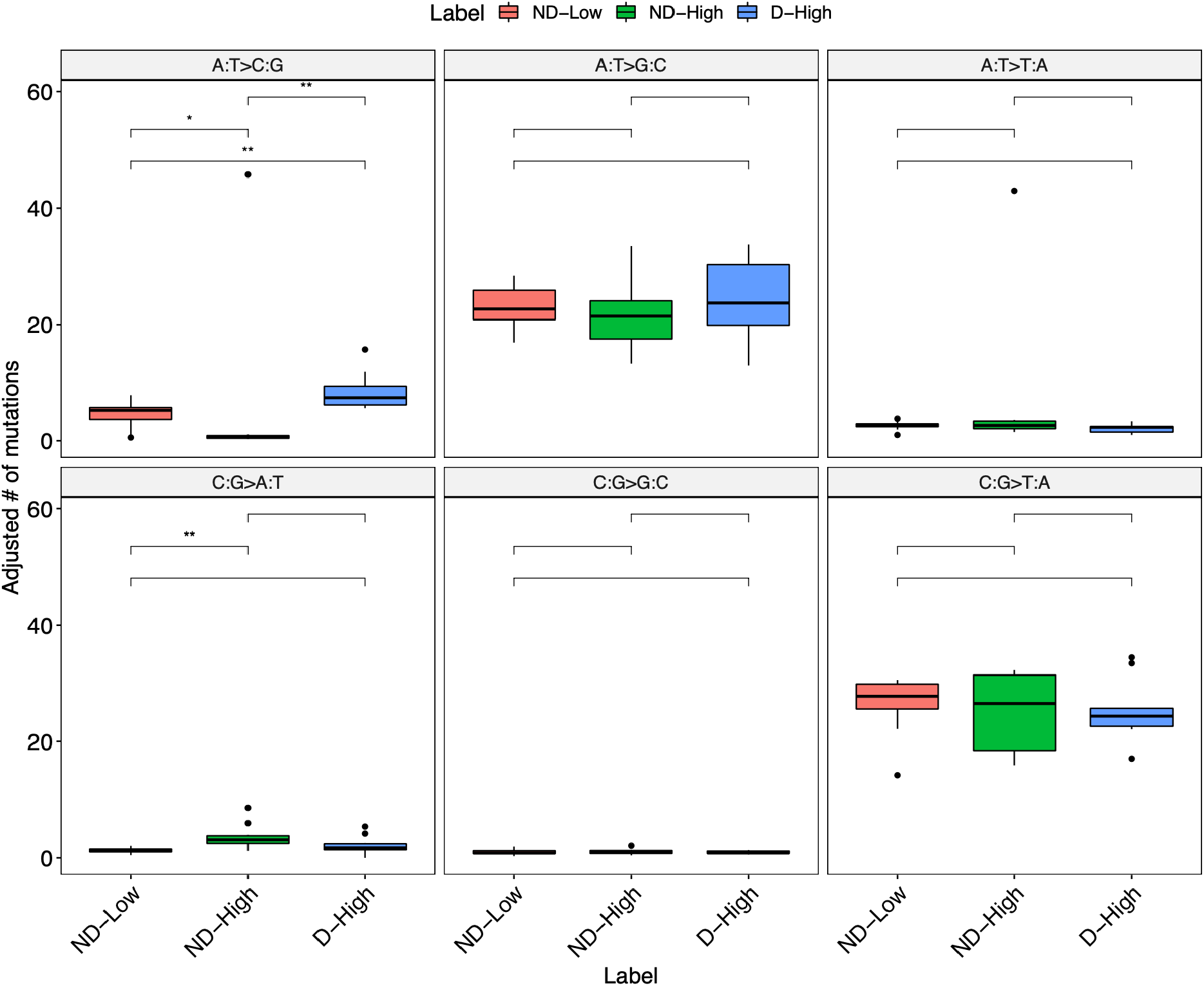
LFMD-revealed mutation spectrum of mtDNA samples with Alzheimer’s Disease (AD) after sensitivity adjustment.

### Ultra-low frequency mutations on *ABL1* associated with potential drug resistance

Using the duplex sequencing method in 2015, Schmitt et al. analysed an individual with chronic myeloid leukaemia who relapsed after targeted therapy with the drug, Imatinib (the Short Read Archive under accession SRR1799908). We analysed this individual and found 5 extra LFMs. Two of them are in the coding region of the *ABL1* gene and change amino acids: E255G and V256G (E, glutamic acid; G, glycine; V, valine). In the drug resistance database of COSMIC[39], we found that 1) E255VDK, change of the 255^th^ amino acid to V, D, or K (D, aspartic acid; K, lysine), is associated with resistance to the drugs Dasatinib, Imatinib, and Nilotinib; and 2) V256L (L, leucine) is related to resistance to the drug Imatinib. Although the amino acid changes, in this case, are not the same as those in the database, these two additional LFMs still infer potential resistance to the drug Imatinib and provide an additional explanation for the clinical relapse of leukemia. Another useful finding is the prediction of potential resistance to Dasatinib and Nilotinib, a result which may guide clinical practice. The annotation results of 5 LFMs are shown in Table 2.

**Table 2.**
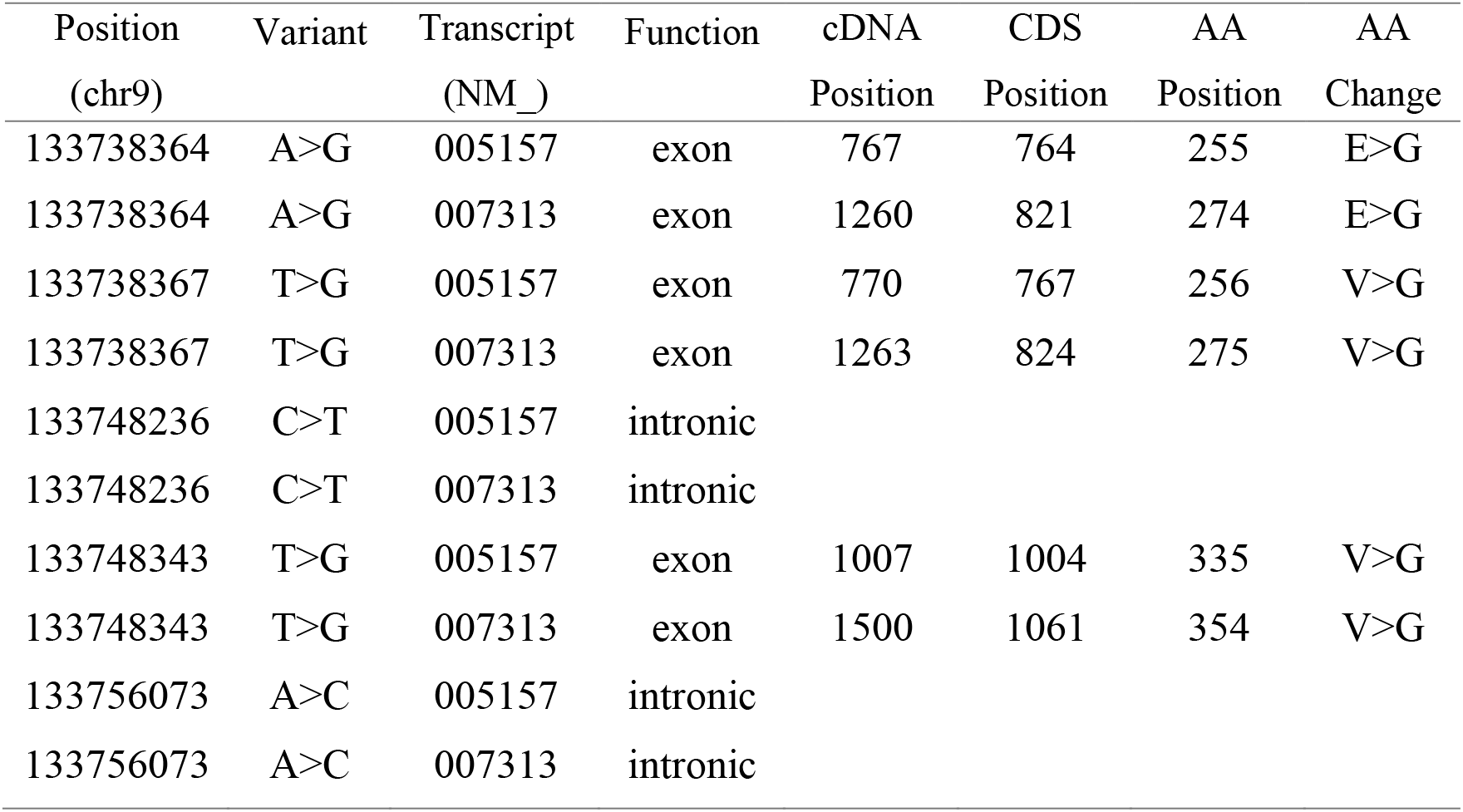
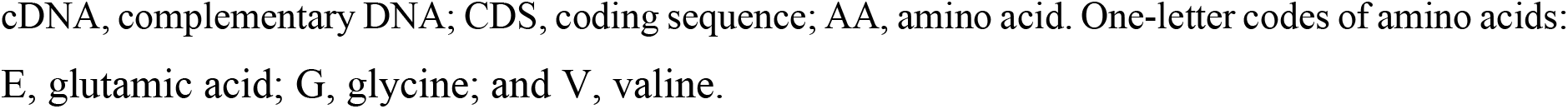
Five low-frequency SNVs on *ABL1* detected by LFMD.

| Position<br>(chr9) | Variant | Transcript<br>(NM_) | Function | cDNA<br>Position | CDS<br>Position | AA<br>Position | AA<br>Change |
| --- | --- | --- | --- | --- | --- | --- | --- |
| 133738364 | A>G | 005157 | exon | 767 | 764 | 255 | E>G |
| 133738364 | A>G | 007313 | exon | 1260 | 821 | 274 | E>G |
| 133738367 | T>G | 005157 | exon | 770 | 767 | 256 | V>G |
| 133738367 | T>G | 007313 | exon | 1263 | 824 | 275 | V>G |
| 133748236 | C>T | 005157 | intronic |  |  |  |  |
| 133748236 | C>T | 007313 | intronic |  |  |  |  |
| 133748343 | T>G | 005157 | exon | 1007 | 1004 | 335 | V>G |
| 133748343 | T>G | 007313 | exon | 1500 | 1061 | 354 | V>G |
| 133756073 | A>C | 005157 | intronic |  |  |  |  |
| 133756073 | A>C | 007313 | intronic |  |  |  |  |

cDNA, complementary DNA; CDS, coding sequence; AA, amino acid. One-letter codes of amino acids: E, glutamic acid; G, glycine; and V, valine.

## Discussion

LFMD is still expensive for target regions >2 Mbp in size because of the need for high sequencing depth. However, as the cost of sequencing continues to fall, it will become more and more practical. In order to sequence a larger region with lower cost, the dilution step of BotSeqS can be introduced into the LFMD pipeline. Because LFMD can deal with tag conflicts, the dilution level can be decreased 1 or 2 magnitudes to increase the sensitivity. Additional experiments will be done soon.

Only accepting randomly sheared DNA fragments, not working on short amplicon sequencing data, and only working on pair-end sequencing data are known limitations of LFMD. Moreover, LFMD’s precision is limited by the accuracy of the alignment software. Although tags are excluded in the model of LFMD described in this paper, LFMD still has the potential to utilize tags and deal with amplicon sequencing data. The basic assumption in our model, that error rates are the same for mutation from the original base to any of the other three bases, is not close enough to reality according to experimental data at present. These remaining issues may be solved in the next version of LFMD.

To estimate the theoretical limit of LFMD, let read length be 100 bp and the standard deviation (SD) of insert size be 20 bp. Furthermore, let N represent the number of read families across one point. Then, N = (2 reads * 100 bases) * (2 directions * 3 SDs * 20 bp) = 24000 if only considering ±3 SD. As the sheering of DNA is not random in the real world, it is safe to set N as 20,000 according to our experience. Ideally, the likelihood ratio test can detect mutations whose frequency is greater than 0.2% in a read family with Q30 bases (refer to Supplementary Figure S2.). Thus, the theoretical limit of minor allele frequency is around 1e-7 (= 0.002 / 20000).

LFMD reduces the cost partly because it discards tags. First, for a typical 100 bp read, the lengths of the tags and the fixed sequences between the tag and the true sequence are 12 bp and 5 bp respectively. So (12+5) / 100 = 17% of data are saved if we do not use tags. Second, the efficiency of targeted capture decreases by about 10 to 20% because of the tags and the fixed sequences, according to in-house experiments. Third, LFMD can work on short read data (PE50) of BGISEQ, and then 30% to 40% of the cost can be saved because of the cheaper sequencing platform. Overall, costs can be reduced by about 70% at best.

Based on the concept of molecular tags, Du Novo[31], UMI-tools[29] and others have made significant improvements since 2012. In contrast, LFMD uses a completely different approach that discards tags in the first place and provides potential advantages. As DS data and high-depth conventional whole genome sequencing data accumulate in public databases, LFMD provides opportunities for further mining of existing datasets, especially mitochondrial DNA.

## Conclusion

To eliminate endogenous tag conflicts, we use a likelihood-based model to separate the read family of the minor allele from that of the major allele. Without additional experimental steps or customized adapters, LFMD achieves higher sensitivity, higher specificity, and lower cost than other methods. LFMD is useful in areas that require ultra-high precision, such as prediction of drug resistance and early screening of cancer. It is a general method that can be used in several cutting-edge areas, and its mathematical methodology can be generalized to increase the power of other NGS tools.

The oxidative damage related mutation, C:G>A:T, is significantly enriched in the mid-stage AD group, which is inconsistent with previous low-frequency mutation studies but consistent with the Mitochondrial Free Radical Theory of Aging. The results suggest that AD may be linked to the aging of brain cells induced by oxidative damage. This success also shows that a tool like LFMD, with better sensitivity and consistency across samples, is essential for further study of disease aetiology.

## Methods and Materials

### Subject recruitment and sampling

A lymphoblastoid cell line (Yan Huang, or YH cell line) established from the first Asian genome donor[40] was used. Total DNA was extracted with the MagPure Buffy Coat DNA Midi KF Kit (MAGEN). The DNA concentration was quantified by Qubit (Invitrogen). DNA integrity was examined by agarose gel electrophoresis. The extracted DNA was kept frozen at −80°C until further processing.

### Mitochondrial genome DNA isolation

Mitochondrial DNA (mtDNA) was isolated and enriched by one or two pairs of primers amplifying the complete mitochondrial genome. The samples were isolated using a single primer set (LR-PCR4) by ultra-high-fidelity Q5 DNA polymerase, following the protocol of the manufacturer (NEB) (Table 3).

**Table 3.**
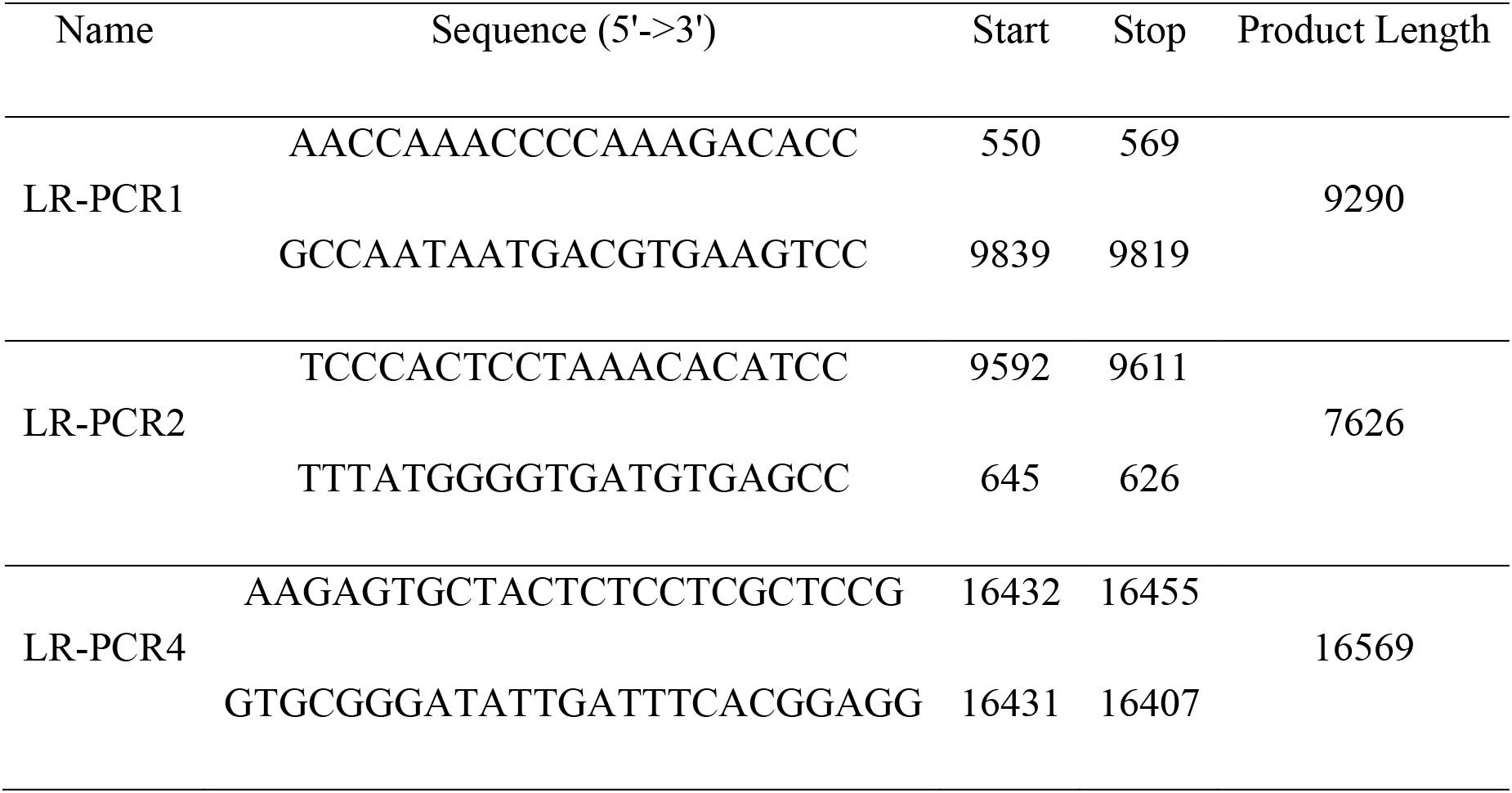
Long-range polymerase chain reaction (LR-PCR) primer sets.

### Library construction and mitochondrial whole genome DNA sequencing

For the BGISeq-500 sequencing platform, mtDNA PCR products were fragmented directly by Covaris E220 (Covaris, Brighton, UK) without purification. Sheared DNA ranging from 150 bp to 500 bp without size selection was purified with an Axygen™ AxyPrep™ Mag PCR Clean-Up Kit. 100 ng of sheared mtDNA was used for library construction. End-repairing and A-tailing was carried out in a reaction containing 0.5 U Klenow Fragment (ENZYMATICS™ P706-500), 6 U T4 DNA polymerase (ENZYMATICS™ P708-1500), 10 U T4 polynucleotide kinase (ENZYMATICS™ Y904-1500), 1 U rTaq DNA polymerase (TAKARA™ R500Z), 5 pmol dNTPs (ENZYMATICS™ N205L), 40 pmol dATPs (ENZYMATICS™ N2010-A-L), 1 X PNK buffer (ENZYMATICS™ B904) and water with a total reaction volume of 50 μl. The reaction mixture was placed in a thermocycler running at 37°C for 30 minutes and heat-denatured at 65°C for 15 minutes with the heated lid at 5°C above the running temperature. Adaptors with 10 bp tags (Ad153-2B) were ligated to the DNA fragments by T4 DNA ligase (ENZYMATICS™ L603-HC-1500) at 25°C. The ligation products were PCR amplified. Twenty to twenty-four purified PCR products were pooled together in equal amounts and then denatured at 95°C and ligated by T4 DNA ligase (ENZYMATICS™ L603-HC-1500) at 37°C to generate a single-strand circular DNA library. Pooled libraries were made into DNA Nanoballs (DNB). Each DNB was loaded into one lane for sequencing.

Sequencing was performed according to the BGISeq-500 protocol (SOP AO) employing the PE100 mode. For reproducibility analyses, YH cell line mtDNA was processed four times following the same protocol as described above to serve as library replicates, and one of the DNBs from the same cell line was sequenced twice as sequencing replicates. A total of 8 datasets were generated using the BGISeq-500 platform. MtDNA sequencing was performed on the BGISeq-500 with 100 bp paired-end reads. The libraries were processed for high-throughput sequencing with a mean depth of ~60000x.

### Method to Generate a Simulated Dataset

We modified Python scripts developed by the Du Novo[31] team to simulate mixed double-strand sequencing data. Although the simulation may not be entirely realistic, the results are still useful to evaluate the performance and the potential drawbacks of LFMD. Of note, the Python scripts use Dr. Heng Li’s well-known program, wgsim, to generate NGS read pairs with controlled sequencing errors, including point errors and indels.

The original script is “sim2.py”, which was developed by the Du Novo team. It can simulate pair-end sequencing with molecular tags. We added a feature that makes the sequencing and PCR error rates slightly higher at both ends of reads than in the middle of reads. The new script is named “sim2_endHighErr.py”. Basically, the error rate at the 5 bp ends of each read is 5 times larger than that in the middle, but not larger than 0.1. This feature makes the simulation data more realistic, although in the real data each base has a different error rate, and bases at read ends do not always have higher error rates.

Apart from calculating the error rate at read ends, we also wrote a script, getFrag_randShear.pl, to randomly shear genome DNA. Because the basic assumption of LFMD is that read pairs from random sheared DNA fragments can be grouped into read families which conduce to reduction of sequencing and PCR error. We set the mean length of sheared fragments as 600 bp and the standard deviation as 20 bp. The lengths of sheared fragments are set to be normally distributed.

With the randomly sheared fragments, we used sim2_endHighErr.py to generate NGS read pairs with original and mutation-inserted mitochondrial reference genomes (NC_012920.1). The base quality in the middle of each read is set as Q20, which means a sequencing error rate of 1%. The PCR error rate is set as 10^−5^, the read length as 100 bp, the length of a molecular tag 12 bp, and the length of the fixed sequence between tag and sequence 5 bp. To be consistent with the data of the Duplex Sequencing system, the fixed sequence is also “TGACT”. In this case, the simulated data can be used as input data for not only LFMD, but also for Duplex Sequencing, UMI-tools, Du Novo, Unified Consensus Maker, etc. Thus, the comparison between different methods can be done on the same dataset. Because LFMD does not need tag information to group reads, we added an extra step to remove tags in the LFMD pipeline to analyze simulated data and Duplex Sequencing data. So, the true read length of data used in LFMD is 83 bp (= 100 - 12 - 5). The results showed that LFMD used 83% of data but outperformed the rival methods which used tags.

The mutation-inserted genome of the mitochondrial chromosome is generated by an in-house script named “fakeMut_rand.pl”. The script accepts parameters like reference genome, numbers of SNVs, number of insertions (INSs), number of deletions (DELs), and maximum lengths of INSs and DELs. Seeds that were used for generating random numbers were included in the output, which is useful because the script can generate exactly the same mutation-inserted genome when the same seed is used. Here is the seed list of the 9 batches: 105901112, 1294553942, 1803264703, 2924365934, 2587769895, 859786530, 2672906100, 3210264384, 4040679464. As the simulation mainly focused on short indels, we set the maximum lengths of indels to 3 bp. We simulated 9 batches of data. Batch 1 has 50 SNVs, 30 INSs, and 30 DELs on a single haplotype of mitochondrial genome. The mutations are distributed randomly, but the minimum distance between two mutations is 7 bp. Batch 2 has 50 SNVs, 20 INSs, and 20 DELs. Batches 3-9 have 100 SNVs, 10 INSs, and 10 DELs. Thus, in 9 batches of data, we simulated a total of 800 SNVs, 120 INSs, and 120 DELs.

Due to the fact that indels may fall next to similar sequences in the genome and result in shifted genome positions in detection results, an additional script, named “blast2mutList.pl”, was used to adjust their genome positions and update their mutation lists. Although most of the shifts can be detected and adjusted, there are several mutations that cannot be dealt with correctly. For example, in the third batch, we found position shifts at 2388 and 11399. These turn out to be issues related to “blastn 2.6.0”. Specifically, the DEL at position 11399, which deletes “TA”, is described falsely by blast as a DEL which deletes “AT” at position 11400. The correct location should be 11399 according to the general standard. Thus, a false positive in the LFMD result may be due to an error in the very beginning induced by the extra tool, “blastn”.

The mutation-inserted mitochondrial genomes were used as templates to generate 15 different numbers of DNA fragments: 100, 200, 300, 400, 500, 600, 700, 800, 900, 1000, 2000, 3000, 4000, 5000, and 10000. Then, based on these fragments, NGS read pairs were generated by simu.randomShear_frag.sh. Similarly, the mitochondrial reference genome without any mutations was also used as template to generate 15 different numbers of DNA fragments: 1e6 - 100, 1e6 - 200, 1e6 - 300, 1e6 - 400, 1e6 - 500, 1e6 - 600, 1e6 - 700, 1e6 - 800, 1e6 - 900, 1e6 - 1000, 1e6 - 2000, 1e6 - 3000, 1e6 - 4000, 1e6 - 5000, and 1e6 - 10000. Thus, after generating NGS read pairs based on these DNA fragments and merging them with read pairs of the corresponding fragments of the mutation-inserted genome, we simulated 15 datasets for each batch. The designed allele frequencies of the 15 datasets are 1e-4, 2e-4, 3e-4, 4e-4, 5e-4, 6e-4, 7e-4, 8e-4, 9e-4, 1e-3, 2e-3, 3e-3, 4e-3, 5e-3, and 1e-2. The overall read depth is around 130,000x for all datasets. Thus, we got simulated sequencing data of 135 (9 batches of data x 15 fragment sets) distinguishable samples.

To evaluate the performance of methods under different sequencing depths, we randomly selected read pairs from these datasets. The fractions of selected read pairs were 0.1, 0.2, 0.25, 0.3, 0.4, 0.5, 0.6, 0.7, 0.8, 0.9, and 1.0. In this way, sequencing data with a total of 11 levels of read depth were generated. The fraction 0.25 corresponded to a read depth of 32,500x, while the fraction 0.5 represented 65,000x.

### Likelihood-based model

We aim to identify alleles at each potential heterozygous position in a read family (grouped according to endogenous tags). Then based on those heterozygous sites, we split the mixed read family into smaller families, and compress each one into a consensus read. Finally, we detect mutations based on all consensus reads, which have much lower error rates than 0.1%.

First, we define a Watson strand as a read pair for which read 1 is the plus strand while read 2 is the minus strand. A Crick strand is defined as a read pair for which read 1 is the minus strand while read 2 is the plus strand. The plus and minus strands are also known as the forward and reverse strands according to the reference genome. Read 1 and 2 are derived from raw pair-end fastq files. Thus, a read family which contains Watson and Crick strand reads simultaneously is an ideal read family because it is supported by both strands of the original DNA template. Second, we select potential heterozygous sites which meet the following criteria: 1) the minor allele is supported by both Watson and Crick reads; 2) minor allele frequencies in both Watson and Crick read family are greater than approximately the average sequencing error rate, often 1% or 0.1%; 3) low-quality bases (<Q20) and low quality alignments (<Q30) are excluded. Finally, we calculate the genotype likelihood in the Watson and Crick family independently in order to eliminate PCR errors during the first PCR cycle.

At each position of a Watson or Crick read family, let *X* denote the sequenced base and *θ* the allele frequencies. Let *P*(*x*|*θ*) be the probability mass function of the random variable *X*, indexed by the parameter *θ* = (*θ*_*A*_, *θ*_*C*_, *θ*_*G*_, *θ*_*T*_)^*T*^, where *θ* belongs to a parameter space Ω. Let *g* ∈ {*A, C, G, T*}, and *θ*_*g*_ represent the frequency of allele *g* at this position. Obviously, we have boundary constraints for *θ*: *θ*_*g*_ ∈ [0, 1] and Σ *θ*_*g*_ = 1.

Assuming *N* reads cover this position, *x*_*i*_ represents the base on read *i* ∈ {1, 2, … , *N*}, and *e*_*i*_ denotes sequencing error of the base, we get

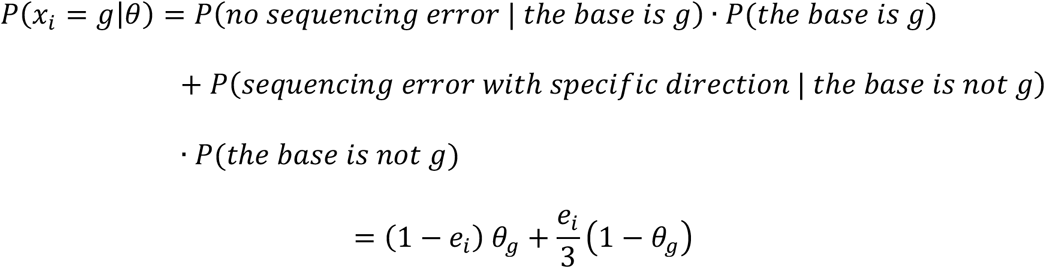

So, the log-likelihood function can be written as

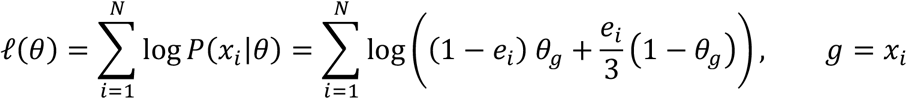

Thus, for each candidate allele *g*, under the null hypothesis *H*_0_: *θ*_*g*_ = 0, *θ* ∈ Ω, and the alternative hypothesis *H*_1_: *θ*_*g*_ ≠ 0, *θ* ∈ Ω, the likelihood ratio test is

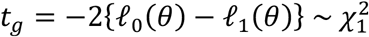

However, as *θ*_*g*_ = 0 lies on the boundary of the parameter space, the general likelihood ratio test needs an adjustment to fit 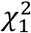. Because the adjustment is related to calculation of a tangent cone[41] in constrained 3-dimensional parameter space, and the computation is too complicated and time-consuming for large scale NGS data, here we use a simplified, straightforward adjustment[42] presented by Chen et al in 2017 (Details in Supplementary Materials).

Interestingly, we finally arrive at a general conclusion that the further adjustment of 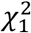 is not helpful in similar cases although the asymptotic distribution we use is not perfect when *N* is small (e.g., N<5). Alternative approaches might be derived in the future. We also compared theoretical P-values with empirical P-values from Monte Carlo procedures (Supplementary Figure S1), explored the power of our model under truly and uniformly distributed sequencing errors (Supplementary Figure S2), and evaluated the accuracy of allele frequency (Supplementary Figure S3 and S4). The simulation results support the theoretical conclusion sufficiently when P-value is equal or less than 10^-3.

Because the null and alternative hypotheses have two and three free variables respectively, the Chi-square distribution has 1 degree of freedom. The P-value of the allele *g* can then be given

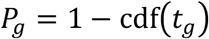

where cdf(*x*) is the cumulative density function of the 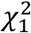 distribution. If *P*_*g*_is less than a given threshold α, the null hypothesis is rejected and the allele *g* is treated as a candidate allele of the read family.

Although *P*_*g*_ cannot be interpreted as the probability that *H*_0,*g*_ is true and allele *g* is an error, it is a proper approximation of the error rate of allele *g* when *P*_*g*_ ≤ 10^−.^. We only reserve alleles with *P*_*g*_ ≤ α in both Watson and Crick families and substitute others with “N”. Then Watson and Crick families are compressed into several single-strand consensus sequences (SSCSs). The SSCSs might contain haplotype information if more than one heterozygous site were detected. Finally, SSCSs which are consistent in both Watson and Crick families are claimed as double-strand consensus sequences (DCSs).

For each allele on a DCS, let *P*_*w*_(*g*) and *P*_*c*_(*g*) represent the relative error rates of the given allele *g* in the Watson and Crick family respectively, and let *P*_*wc*_(*g*) denote the error rate of the allele *g* on the DCS. Thus,

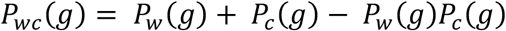

For a read family which proliferated from ***n*** original templates, a coalescent model can be used to model the PCR procedure[43]. The exact coalescent PCR error rate is too complicated to be calculated quickly, so we tried to give a rough estimate. According to the model, a PCR error proliferates, and its frequency decreases exponentially with the number of PCR cycles, ***m***. For example, an error that occurs in the first PCR cycle would occupy half of the PCR products, an error that occurs in the second cycle occupies a quarter, the third only 1/8, and so on. As we only need to consider PCR errors which are detectable, the coalescent PCR error rate is defined as the probability to detect a PCR error whose frequency ≥ 2^−***m***^/***n***, and it is equal to or less than

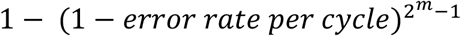

Let *e*_*pcr*_(*g*) denote the coalescent PCR error rate and *P*_*pcr*_(*g*) the united PCR error rate of the double strand consensus allele. Then we get

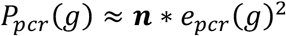

Because the PCR fidelity ranges from 10^−5^ to 10^−7^, we get *P*_*wc*_(*g*)*P*_*pcr*_(*g*) ≈ 0, then the combined base quality of the allele on the DCS is

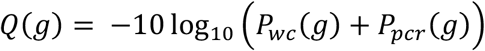

Then *Q*(*g*) is transferred to an ASCII character, and a series of characters make a base quality sequence for the DCS. Finally, we generate a BAM file with DCSs and their quality sequences.

With the BAM file which contains all the high-quality DCS reads, the same approach is used to give each allele a P-value at each genomic position which is covered by DCS reads. Adjusted P-values (q-values) are given via the Benjamin-Hochberg procedure. The threshold of q-values is selected according to the total number of tests conducted and false discovery rate (FDR) which can be accepted.

A similar mathematical model was described in detail in previous papers by Ding et al[6] and Guo et al[20]. Ding et al. used this model to reliably call mutations with frequency > 4%. In contrast, we use this model to deal with read families rather than non-duplicate reads. In a mixed read family, most of the minor allele frequencies are larger than 4%, so the power of the model meets our expectation.

For those reads containing indels, the CIGAR strings in BAM files contain I or D. It is obvious that reads with different CIGAR strings cannot fit into one read family. Thus, CIGAR strings can also be used as part of endogenous tags. In contrast, the soft-clipped part of CIGAR strings cannot be ignored when considering start and end positions because low-quality parts of reads tend to be clipped, and the coordinates after clipping are not a proper endogenous tag for the original DNA template.

### Method to estimate sensitivity

One of the major concerns of low frequency mutation detection is the relatively-low sensitivity and high-depth requirement due to the fact that the sensitivity of LFMD might be affected by total sequencing depth, depth distribution, randomness of DNA shearing, distribution of length of sheared DNA fragment, DCS depth, PCR bias, number of PCR cycles during library construction, initial amount of DNA, GC content, etc. Thus, comparison among multiple samples with various library construction and sequencing status is challenging (Supplementary Figure S5). To overcome this obstacle, it is essential to carry out a robust sensitivity estimation for all samples before comparing their mutation calling results. With the estimated sensitivity, the number of called mutations can be normalized into a pseudo-number of mutations, then the comparison between the samples will be fair (Supplementary Figure S6). In addition, accurate estimation of the number of fragments in a read family can help to estimate the PCR error rate accurately and thus improve the specificity of LFMD.

As shown in Figure 6, insert size is defined as the length of DNA fragment after random shearing in the library construction step; bin is defined as a possible genome position that can be occupied by a DNA fragment with given insert size. As the read length is given and fixed, the total number of bins are determined and fixed. The positions that are not occupied are empty bins.

**Figure 6.**
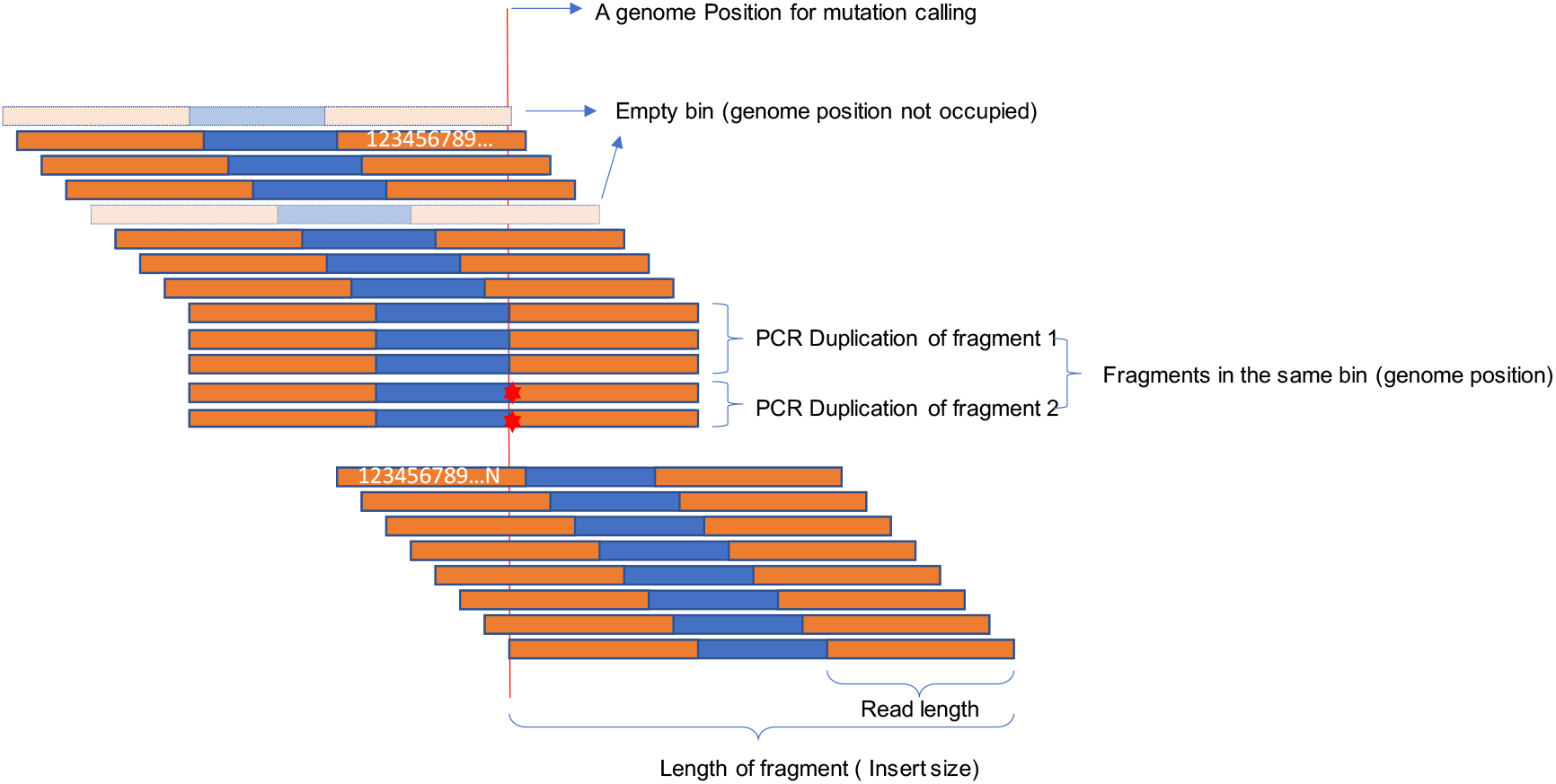
The definition of insert size and bin in the LFMD context.

For each DNA fragment, let *Y* denote the number of sequenced read pairs of the fragment. Assume *Y* is a Poisson distributed random variable with mean *λ*_2_ and variance *λ*_2_, denoted as *Y*~*Pois*(*λ*_2_). So, its probability mass function can be written as

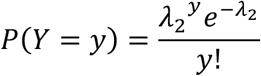

For *k* fragments in a bin,

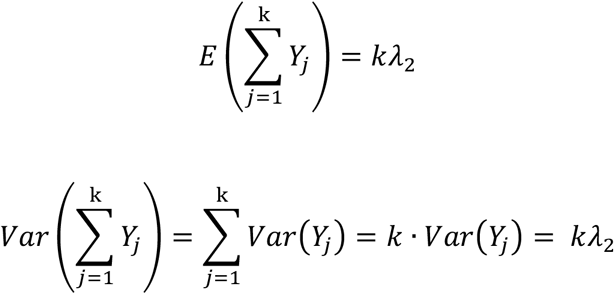

Let 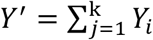 and *λ*′ = *kλ*_2_, then *Y*′~*Pois*(*λ*′), where *Y*′ is the number of sequenced read pairs in a bin which has *k* fragments. Let *X* denote the number of fragments in a bin. Assume *X* follows a Poisson distribution, *X*~*Pois*(*λ*), with mean *λ* and variance *λ*.

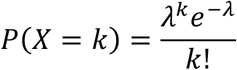

So, the probability that *y* read pairs found in a bin is

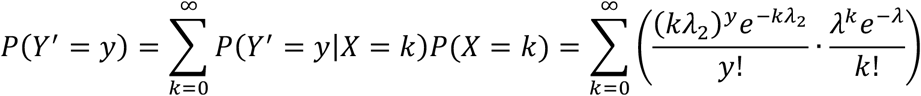

Let *N*_*b*_ be the number of bins, and *Z* be the total number of all read pairs in all bins, then

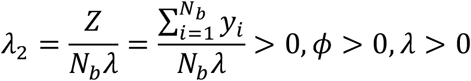

Let *N*_*empty*_ denote the number of empty bins, then *P*(*Y*′ = 0) ≈ *N*_*empty*_/*N*_*b*_. Therefore, *λ* can be calculated via solving the following equation:

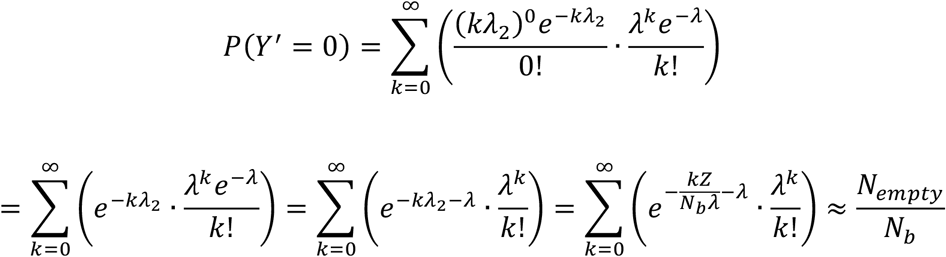

However, the equation, 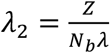 and *P*(*Y*′ = 0) ≈ *N*_*empty*_/*N*_*b*_, may not hold when the amount of data is small. So, the simple estimation of λ is efficient but biased.

To estimate *λ* and *λ*_2_ generally, a better method is to select them to maximize the log-likelihood function

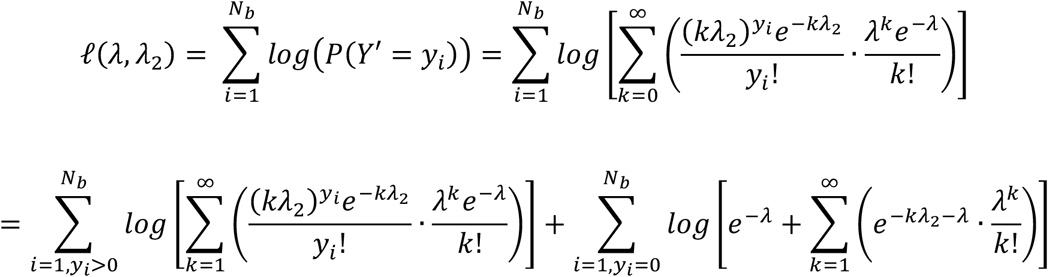

Once the Maximum Likelihood Estimators (MLEs) are obtained for the log-likelihood function, it is possible to calculate the probability of *k* fragments in a bin given observed *y* read pairs according to Bayes’ theorem:

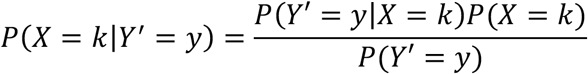

We assume only one fragment in the bin has the mutation, considering the low frequency of the mutation. Thus, in the bin, the mutation frequency is 1/*k*. Let *s* denote the number of reads supporting the mutation. Given *y* reads and the threshold that at least 3 reads support the mutation, the probability that the mutation cannot be detected is

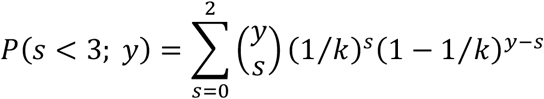

If considering the sequencing error as *e*, it is possible to add the probability that 3 PCR products of the fragment support the mutation but at least one of them has a sequencing error, wherein the mutation cannot be detected:

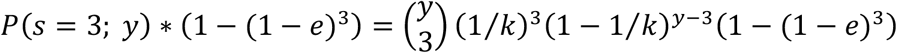

Let *y*_*w*_ and *y*_*c*_ denote the number of read pairs in Watson and Crick families respectively. We get the probability that the mutation cannot be detected as

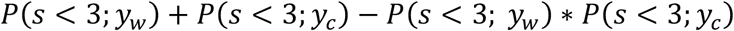

So, we can estimate the sensitivity at the position as

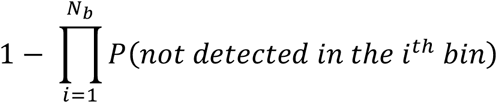

To fit the empirical sensitivities better, more sophisticated models, including overdispersion of the Poisson Model and stand bias, were considered in the supplementary materials. In fact, the simple two-Poisson model is good enough to fit the sensitivity curve, especially when sequencing depth is low (<60,000x).

## Declarations

### Ethics approval and consent to participate

Not applicable.

### Consent for publication

All authors have approved the manuscript for submission.

### Availability of data and materials

The datasets supporting the conclusions of this article are available in the CNGBdb repository (https://db.cngb.org/cnsa/) with accession code CNP0000297. Analysis codes used in this paper can be accessed at https://github.com/RainyEricYe/LFMD.

### Authors’ contributions

P.C.S. and C.N. supervised the entire work. R.Y. wrote the manuscript. R.Y. and X.H.Z. wrote the code and evaluated the performance of LFMD. Y.W.Q. assisted the analysis of mitochondrial data. R.J. led all wet-lab experiments and tested the LR-PCR on mitochondria. R.J., Y.T.A., and J.M.X. prepared the samples and did the experiments. T.M. and P.C.S. helped the mathematical part. P.C.S, C.N., L.B., X.X., H.M.Y, X.Q.Z, X.L., and T.M. contributed to critical revision of the manuscript. All authors read and approved the final manuscript.

## Acknowledgements

We thank Prof. Scott R. Kennedy for kindly sharing their valuable data. We thank Jeremiah Wala and Rameen Beroukhim for developing a wonderful C++ API and SeqLib[44], and we thank S. Bochkanov, V. Bystritsky, and other contributors for the development and maintenance of ALGLIB[45], which accelerated the coding progress of LFMD significantly. We also thank Heng Li, Richard Durbin and other developers for creating bwa[46] and samtools[47]. We are so grateful to McKenna et al. and Broad Institute for the introduction and maintenance of the GATK[48] framework. And we do appreciate the open source scripts of Nicholas Stoler et al. to simulate tagged sequences[31]. Without the open source spirit and knowledge sharing, our work could not be done.

## References

1. Newman AM, Lovejoy AF, Klass DM, Kurtz DM, Chabon JJ, Scherer F, Stehr H, Liu CL, Bratman SV, Say C: Integrated digital error suppression for improved detection of circulating tumor DNA. Nature biotechnology 2016, 34:547.

2. Phallen J, Sausen M, Adleff V, Leal A, Hruban C, White J, Anagnostou V, Fiksel J, Cristiano S, Papp E: Direct detection of early-stage cancers using circulating tumor DNA. Science translational medicine 2017, 9:eaan2415.

3. Newman AM, Bratman SV, To J, Wynne JF, Eclov NC, Modlin LA, Liu CL, Neal JW, Wakelee HA, Merritt RE: An ultrasensitive method for quantitating circulating tumor DNA with broad patient coverage. Nature medicine 2014, 20:548.

4. Hoekstra JG, Hipp MJ, Montine TJ, Kennedy SR: Mitochondrial DNA mutations increase in early stage Alzheimer disease and are inconsistent with oxidative damage. Annals of neurology 2016, 80:301–306.

5. Simon DK, Lin MT, Zheng L, Liu G-J, Ahn CH, Kim LM, Mauck WM, Twu F, Beal MF, Johns DR: Somatic mitochondrial DNA mutations in cortex and substantia nigra in aging and Parkinson’s disease. Neurobiology of aging 2004, 25:71–81.

6. Ding J, Sidore C, Butler TJ, Wing MK, Qian Y, Meirelles O, Busonero F, Tsoi LC, Maschio A, Angius A: Assessing mitochondrial DNA variation and copy number in lymphocytes of~ 2,000 Sardinians using tailored sequencing analysis tools. PLoS genetics 2015, 11:e1005306.

7. Wallace DC, Chalkia D: Mitochondrial DNA genetics and the heteroplasmy conundrum in evolution and disease. Cold Spring Harbor perspectives in biology 2013, 5:a021220.

8. Schmitt MW, Fox EJ, Prindle MJ, Reid-Bayliss KS, True LD, Radich JP, Loeb LA: Sequencing small genomic targets with high efficiency and extreme accuracy. Nature methods 2015, 12:423.

9. Dimond R: Social and ethical issues in mitochondrial donation. British medical bulletin 2015, 115:173.

10. Jabara CB, Jones CD, Roach J, Anderson JA, Swanstrom R: Accurate sampling and deep sequencing of the HIV-1 protease gene using a Primer ID. Proceedings of the National Academy of Sciences 2011, 108:20166–20171.

11. Marquis J, Lefebvre G, Kourmpetis YA, Kassam M, Ronga F, De Marchi U, Wiederkehr A, Descombes P: MitoRS, a method for high throughput, sensitive, and accurate detection of mitochondrial DNA heteroplasmy. BMC genomics 2017, 18:326.

12. Kang E, Wang X, Tippner-Hedges R, Ma H, Folmes CD, Gutierrez NM, Lee Y, Van Dyken C, Ahmed R, Li Y: Age-related accumulation of somatic mitochondrial DNA mutations in adult-derived human iPSCs. Cell Stem Cell 2016, 18:625–636.

13. Blandini F, Greenamyre JT, Nappi G: The role of glutamate in the pathophysiology of Parkinson’s disease. Functional neurology 1996, 11:3–15.

14. Navin N, Kendall J, Troge J, Andrews P, Rodgers L, McIndoo J, Cook K, Stepansky A, Levy D, Esposito D: Tumour evolution inferred by single-cell sequencing. Nature 2011, 472:90.

15. Baslan T, Hicks J: Single cell sequencing approaches for complex biological systems. Current opinion in genetics & development 2014, 26:59–65.

16. Lou DI, Hussmann JA, McBee RM, Acevedo A, Andino R, Press WH, Sawyer SL: High-throughput DNA sequencing errors are reduced by orders of magnitude using circle sequencing. Proceedings of the National Academy of Sciences 2013, 110:19872–19877.

17. Wang K, Lai S, Yang X, Zhu T, Lu X, Wu C-I, Ruan J: Ultrasensitive and high-efficiency screen of de novo low-frequency mutations by o2n-seq. Nature communications 2017, 8:15335.

18. Kinde I, Wu J, Papadopoulos N, Kinzler KW, Vogelstein B: Detection and quantification of rare mutations with massively parallel sequencing. Proceedings of the National Academy of Sciences 2011, 108:9530–9535.

19. Vogelstein B, Kinzler KW, Papadopoulos N, Kinde I: Safe sequencing system. Google Patents; 2016.

20. Guo Y, Li J, Li C-I, Shyr Y, Samuels DC: MitoSeek: extracting mitochondria information and performing high-throughput mitochondria sequencing analysis. Bioinformatics 2013, 29:1210–1211.

21. Kim J, Kim D, Lim JS, Maeng JH, Son H, Kang H-C, Nam H, Lee JH, Kim S: The use of technical replication for detection of low-level somatic mutations in next-generation sequencing. Nature communications 2019, 10:1047.

22. Wang TT, Abelson S, Zou J, Li T, Zhao Z, Dick JE, Shlush LI, Pugh TJ, Bratman SV: High efficiency error suppression for accurate detection of low-frequency variants. Nucleic acids research 2019.

23. Hoang ML, Kinde I, Tomasetti C, McMahon KW, Rosenquist TA, Grollman AP, Kinzler KW, Vogelstein B, Papadopoulos N: Genome-wide quantification of rare somatic mutations in normal human tissues using massively parallel sequencing. Proceedings of the National Academy of Sciences 2016, 113:9846–9851.

24. Schmitt MW, Kennedy SR, Salk JJ, Fox EJ, Hiatt JB, Loeb LA: Detection of ultra-rare mutations by next-generation sequencing. Proceedings of the National Academy of Sciences 2012, 109:14508–14513.

25. Kennedy SR, Schmitt MW, Fox EJ, Kohrn BF, Salk JJ, Ahn EH, Prindle MJ, Kuong KJ, Shen JC, Risques RA, Loeb LA: Detecting ultralow-frequency mutations by Duplex Sequencing. Nat Protoc 2014, 9:2586–2606.

26. Wang K, Lai S, Yang X, Zhu T, Lu X, Wu C-I, Ruan J: Ultrasensitive and high-efficiency screen of de novo low-frequency mutations by o2n-seq. Nature communications 2017, 8:1–11.

27. Peng Q, Satya RV, Lewis M, Randad P, Wang Y: Reducing amplification artifacts in high multiplex amplicon sequencing by using molecular barcodes. BMC genomics 2015, 16:589.

28. Kou R, Lam H, Duan H, Ye L, Jongkam N, Chen W, Zhang S, Li S: Benefits and challenges with applying unique molecular identifiers in next generation sequencing to detect low frequency mutations. PloS one 2016, 11:e0146638.

29. Smith T, Heger A, Sudbery I: UMI-tools: modeling sequencing errors in Unique Molecular Identifiers to improve quantification accuracy. Genome Res 2017, 27:491–499.

30. Wilm A, Aw PPK, Bertrand D, Yeo GHT, Ong SH, Wong CH, Khor CC, Petric R, Hibberd ML, Nagarajan N: LoFreq: a sequence-quality aware, ultra-sensitive variant caller for uncovering cell-population heterogeneity from high-throughput sequencing datasets. Nucleic Acids Research 2012, 40:11189–11201.

31. Stoler N, Arbeithuber B, Guiblet W, Makova KD, Nekrutenko A: Streamlined analysis of duplex sequencing data with Du Novo. Genome biology 2016, 17:180.

32. Braak H, Braak E: Neuropathological stageing of Alzheimer-related changes. Acta neuropathologica 1991, 82:239–259.

33. Cheng KC, Cahill DS, Kasai H, Nishimura S, Loeb LA: 8-Hydroxyguanine, an abundant form of oxidative DNA damage, causes G----T and A C substitutions. Journal of Biological Chemistry 1992, 267:166–172.

34. Cooke MS, Evans MD, Dizdaroglu M, Lunec J: Oxidative DNA damage: mechanisms, mutation, and disease. The FASEB Journal 2003, 17:1195–1214.

35. Marnett LJ: Oxyradicals and DNA damage. carcinogenesis 2000, 21:361–370.

36. Lovell MA, Soman S, Bradley MA: Oxidatively modified nucleic acids in preclinical Alzheimer’s disease (PCAD) brain. Mechanisms of ageing and development 2011, 132:443–448.

37. Wang J, Xiong S, Xie C, Markesbery W, Lovell M: Increased oxidative damage in nuclear and mitochondrial DNA in Alzheimer’s disease. Journal of neurochemistry 2005, 93:953–962.

38. Wang J, Markesbery WR, Lovell MA: Increased oxidative damage in nuclear and mitochondrial DNA in mild cognitive impairment. Journal of neurochemistry 2006, 96:825–832.

39. Tate JG, Bamford S, Jubb HC, Sondka Z, Beare DM, Bindal N, Boutselakis H, Cole CG, Creatore C, Dawson E: COSMIC: the catalogue of somatic mutations in cancer. Nucleic acids research 2018, 47:D941–D947.

40. Wang J, Wang W, Li R, Li Y, Tian G, Goodman L, Fan W, Zhang J, Li J, Zhang J, et al: The diploid genome sequence of an Asian individual. Nature 2008, 456:60–65.

41. Drton M: Likelihood ratio tests and singularities. The Annals of Statistics 2009, 37:979–1012.

42. Chen Y, Huang J, Ning Y, Liang K-Y, Lindsay BG: A conditional composite likelihood ratio test with boundary constraints. Biometrika 2017, 105:225–232.

43. Weiss G, Von Haeseler A: A coalescent approach to the polymerase chain reaction. Nucleic acids research 1997, 25:3082–3087.

44. Wala J, Beroukhim R: SeqLib: a C++ API for rapid BAM manipulation, sequence alignment and sequence assembly. Bioinformatics 2016, 33:751–753.

45. Bochkanov S, Bystritsky V: Alglib. Available from: www.alglib.net 2013, 59.

46. Li H, Durbin R: Fast and accurate short read alignment with Burrows–Wheeler transform. Bioinformatics 2009, 25:1754–1760.

47. Li H, Handsaker B, Wysoker A, Fennell T, Ruan J, Homer N, Marth G, Abecasis G, Durbin R: The sequence alignment/map format and SAMtools. Bioinformatics 2009, 25:2078–2079.

48. McKenna A, Hanna M, Banks E, Sivachenko A, Cibulskis K, Kernytsky A, Garimella K, Altshuler D, Gabriel S, Daly M: The Genome Analysis Toolkit: a MapReduce framework for analyzing next-generation DNA sequencing data. Genome research 2010, 20:1297–1303.

