## Supplementary material for "LFMD: detecting low-frequency mutations in high-depth genome sequencing data without molecular tags": sup

### Supplementary Materials

#### Adjustment at the boundary of parameter space

In order to utilize Chen et al.'s method[1], we need to introduce conditional events. Let $\{\mathcal{A}_1, \dots, \mathcal{A}_K\}$ denote the set of conditional events which are mapped to four alleles at the position. Let $k = 1, \dots, K, and\ K = 4$ be the total number of events. We get the log-likelihood component

$$\ell_k\{\theta;\ \mathcal{A}_k(x_i)\} = \sum\nolimits_{x_i \in \mathcal{A}_k} \log P(x_i|\theta)$$

Then the composite conditional log-likelihood can be constructed as

$$\ell_c(\theta) = \sum_{i=1}^{N} \sum_{k=1}^{K} \omega_{ik} \ell_k\{\theta;\ \mathcal{A}(x_i)\}$$

in which we set

$$\omega_{ik} = 1$$

Let $\hat{\theta}_c = \arg\max_{\theta \in \Omega} \ell_c(\theta)$ be the maximum composite likelihood estimator, and define the composite score function, sensitivity matrix, and variability matrix respectively as

$$U_c(\theta) = \frac{\partial \ell_c(\theta)}{\partial \theta}$$

$$H = \lim_{N \to \infty} -\frac{1}{N}\, E\left\{\frac{\partial^2 \ell_c(\theta)}{\partial \theta^T \partial \theta}\right\}$$

$$V = \lim_{N \to \infty} \frac{1}{N} E\left[\left\{\frac{\partial \ell_c(\theta)}{\partial \theta}\right\}\left\{\frac{\partial \ell_c(\theta)}{\partial \theta}\right\}^T\right]$$

The corresponding estimators of $H$ and $V$ are denoted by $\hat{H}$ and $\hat{V}$ evaluated at $\hat{\theta}_c$. The modified composite likelihood under boundary constraints was given by Chen et al[1] as

$$\ell_M(\theta) = \ell_c(\hat{\theta}_c) - \{T(\theta)^T \hat{H}_A T(\theta)\}\phi(\theta)$$

where

$$T(\theta) = N^{-1/2}\hat{H}^{-1}U_c(\hat{\theta}_c) - N^{1/2}(\theta - \hat{\theta}_c)$$

$$\hat{H}_A = \hat{H}\hat{V}^{-1}\hat{H}$$

$$\phi(\theta) = \frac{\ell_c(\theta) - \ell_c(\hat{\theta}_c)}{-T(\theta)^T\hat{H}T(\theta) + N^{-1}U_c(\hat{\theta}_c)^T\hat{H}^{-1}U_c(\hat{\theta}_c)}$$

Thus, we derive the adjusted likelihood ratio test

$$t_g = -2\{\ell_M(\theta_0) - \ell_M(\hat{\theta}_M)\} \sim \chi_1^2$$

where $\hat{\theta}_M = \arg\max_{\theta\in\Omega} \ell_{\mathrm{M}}(\theta)$ and $\theta_0$ is the parameter $\theta$ under null hypothesis $H_0$.

To facilitate the calculation of $H$ and $V$, we let $pmf(e)$ denote the probability mass function of sequencing error rate $e$, and the expected number of bases with $e$ is represented as

$$\lim_{N\to\infty} N \cdot pmf(e)$$

Then, the expected number of bases $g$ with $e$ is

$$\lim_{N\to\infty} N \cdot pmf(e) \cdot \left\{(1-e)\,\theta_g + \frac{e}{3}(1-\theta_g)\right\}$$

Thus,

$$E\left[\frac{\partial \ell_c(\theta)}{\partial \theta_g}\right] = \lim_{N\to\infty} \sum\nolimits_e \left\{N \cdot pmf(e) \cdot \left\{(1-e)\,\theta_g + \frac{e}{3}(1-\theta_g)\right\} \cdot \frac{1-\frac{4e}{3}}{(1-e)\,\theta_g + \frac{e}{3}(1-\theta_g)}\right\}$$

$$= \lim_{N\to\infty} N \cdot \sum\nolimits_e \left\{pmf(e)\left(1 - \frac{4e}{3}\right)\right\} = \lim_{N\to\infty} N \cdot C$$

where $C$ is a finite constant. Then we derive

$$V = \lim_{N\to\infty} \frac{1}{N} E\left[\left\{\frac{\partial \ell_c(\theta)}{\partial \theta}\right\}\left\{\frac{\partial \ell_c(\theta)}{\partial \theta}\right\}^T\right] = \lim_{N\to\infty} NC^2 \begin{Bmatrix} 1 & 1 & 1 & 1 \\ 1 & 1 & 1 & 1 \\ 1 & 1 & 1 & 1 \\ 1 & 1 & 1 & 1 \end{Bmatrix}$$

As a result, $\hat{V}^{-1}$ tends to $\boldsymbol{0}$ in the model, which means the adjustment is not necessary. To be clear, the special form of matrix $V$ with all equal elements is due to the infinite $N$, which ensures all possible $e$ and $g$ occur in the function $\ell_c(\theta)$. $V$ does not have a special form when $N$ is a finite number. The simulation results are concordant with the theoretical results (Figure S1). The power of the model is evaluated in Figure S2.

**Two-Poisson model with over-dispersion where the over-dispersion parameter is for the second Poisson distribution**

For each DNA fragment, let $Y$ denote the number of sequenced read pairs of the fragment. Assume $Y$ is a Negative Binomial distributed random variable with mean $\mu$ and dispersion $\phi$, denoted as $Y \sim NB(\mu, \phi)$. We choose the parameterization such that the probability mass function is

$$f(y; \mu, \phi) = P(Y = y) = \frac{\Gamma(y + \phi^{-1})}{\Gamma(\phi^{-1})\Gamma(y + 1)} \left(\frac{1}{1 + \mu\phi}\right)^{\phi^{-1}} \left(\frac{\mu}{\phi^{-1} + \mu}\right)^{y}$$

giving $E(Y) = \mu$ and $Var(Y) = \mu + \phi\mu^2$. We focus here on over-dispersion where $\phi > 0$, though $\phi > -\mu^{-1}$ is in fact permitted by the model. Actually, $\phi < 0$ means under-dispersion rather than over-dispersion.

For $k$ fragments in a bin,

$$E\left(\sum_{j=1}^{k} Y_j\right) = k\mu$$

$$Var\left(\sum_{j=1}^{k} Y_j\right) = \sum_{j=1}^{k} Var(Y_j) = k \cdot Var(Y_j) = k\mu + \frac{\phi}{k}(k\mu)^2$$

Let $Y' = \sum_{j=1}^{\mathrm{k}} Y_i$ , $\mu' = k\mu$ , and $\phi' = \frac{\phi}{k}$ . $Y' \sim NB(k\mu,\ \frac{\phi}{k})$ for the bin which has $k$ fragments. Let $X$ denote the number of fragments in a bin. $X$ follows a Poisson distribution $X \sim Pois(\lambda)$ with mean $\lambda$ and variance $\lambda$.

$$P(X = k) = \frac{\lambda^k e^{-\lambda}}{k!}$$

So, the probability that $y$ read pairs are found in a bin is

$$P(Y' = y) = \sum_{k=0}^{\infty} P(Y' = y | X = k) P(X = k)$$

$$= e^{-\lambda} f(y) + \sum_{k=1}^{\infty} P(Y' = y | X = k) P(X = k)$$

$$= e^{-\lambda} f(y) + \sum_{k=1}^{\infty} \left\{ \frac{\Gamma(y + {\phi'}^{-1})}{\Gamma({\phi'}^{-1})\Gamma(y+1)} \left(\frac{1}{1 + \mu'\phi'}\right)^{{\phi'}^{-1}} \left(\frac{\mu'}{{\phi'}^{-1} + \mu'}\right)^{y} \cdot \frac{\lambda^k e^{-\lambda}}{k!} \right\}$$

$$= e^{-\lambda} f(y) + \sum_{k=1}^{\infty} \left\{ \frac{\Gamma\left(y + \frac{k}{\phi}\right)}{\Gamma\left(\frac{k}{\phi}\right)\Gamma(y+1)} \left(\frac{1}{1 + k\mu\frac{\phi}{k}}\right)^{\frac{k}{\phi}} \left(\frac{k\mu}{\frac{k}{\phi} + k\mu}\right)^{y} \cdot \frac{\lambda^k e^{-\lambda}}{k!} \right\}$$

$$= e^{-\lambda} f(y) + \sum_{k=1}^{\infty} \left\{ \frac{\Gamma\left(y + \frac{k}{\phi}\right)}{\Gamma\left(\frac{k}{\phi}\right)\Gamma(y+1)} \left(\frac{1}{1 + \mu\phi}\right)^{\frac{k}{\phi}} \left(\frac{1}{1 + (\mu\phi)^{-1}}\right)^{y} \cdot \frac{\lambda^k e^{-\lambda}}{k!} \right\}$$

in which

$$f(y) = \begin{cases} 1, & y = 0 \\ 0, & y > 0 \end{cases}$$

Given $N_b$ is the number of bins, and $Z$ is the total number of all read pairs in all bins, then

$$\mu = \frac{Z}{N_b\lambda} = \frac{\sum_{i=1}^{N_b} y_i}{N_b\lambda} > 0, \phi > 0, \lambda > 0$$

Select $\mu$ and $\phi$ to maximize log-likelihood function

$$\ell(\mu, \phi) = \sum_{i=1}^{N_b} log\big(P(Y' = y_i)\big)$$

$$= \sum_{i=1}^{N_b} log\left\{e^{-\lambda} f(y_i) + \sum_{k=1}^{\infty}\left\{\frac{\Gamma\left(y_i + \frac{k}{\phi}\right)}{\Gamma\left(\frac{k}{\phi}\right)\Gamma(y_i + 1)}\left(\frac{1}{1 + \mu\phi}\right)^{\frac{k}{\phi}}\left(\frac{1}{1 + (\mu\phi)^{-1}}\right)^{y_i} \cdot \frac{\lambda^k e^{-\lambda}}{k!}\right\}\right\}$$

$$= \sum_{i=1, y_i>0}^{N_b} log\left\{\sum_{k=1}^{\infty}\left\{\frac{\Gamma\left(y_i + \frac{k}{\phi}\right)}{\Gamma\left(\frac{k}{\phi}\right)\Gamma(y_i + 1)}\left(\frac{1}{1 + \mu\phi}\right)^{\frac{k}{\phi}}\left(\frac{1}{1 + (\mu\phi)^{-1}}\right)^{y_i} \cdot \frac{\lambda^k e^{-\lambda}}{k!}\right\}\right\}$$

$$+ \sum_{i=1, y_i=0}^{N_b} log\left\{e^{-\lambda} + \sum_{k=1}^{\infty}\left\{\left(\frac{1}{1 + \mu\phi}\right)^{\frac{k}{\phi}} \cdot \frac{\lambda^k e^{-\lambda}}{k!}\right\}\right\}$$

**Two-Poisson Model with over-dispersion where the over-dispersion parameter is for the first Poisson distribution**

Let $X$ denote the number of fragments in a bin. $X$ is a negative binomial distributed random variable with mean $\mu$ and dispersion $\phi$, which is denoted as $X \sim NB(\mu, \phi)$. We choose the parameterization such that the probability mass function is

$$f(x; \mu, \phi) = P(X = x) = \frac{\Gamma(x + \phi^{-1})}{\Gamma(\phi^{-1})\Gamma(x + 1)}\left(\frac{1}{1 + \mu\phi}\right)^{\phi^{-1}}\left(\frac{\mu}{\phi^{-1} + \mu}\right)^{x}$$

giving $E(X) = \mu$ and $Var(X) = \mu + \phi\mu^2$. We focus here on over-dispersion where $\phi > 0$, though $\phi > -\mu^{-1}$ is in fact permitted by the model.

Let $Y$ denote the number of sequenced read pairs per fragment. $Y$ follows a Poisson distribution $Y \sim Pois(\lambda)$ with mean $\lambda$ and variance $\lambda$.

$$P(Y = y) = \frac{\lambda^y e^{-\lambda}}{y!}$$

For $k$ fragments in a bin,

$$E\left(\sum_{j=1}^{\mathrm{x}} Y_j\right) = x\lambda$$

$$Var\left(\sum_{j=1}^{\mathrm{x}} Y_j\right) = \sum_{j=1}^{\mathrm{x}} Var(Y_j) = x \cdot Var(Y_j) = x\lambda$$

Let $Y' = \sum_{j=1}^{\mathrm{k}} Y_i$ and $\lambda' = x\lambda$. We have $Y' \sim Pois(\lambda')$, where $Y'$ is the number of sequenced read pairs in a bin with $x$ fragments.

So, the probability that $y$ read pairs are found in a bin is

$$P(Y' = y) = \sum_{x=0}^{\infty} P(Y' = y | X = x) P(X = x)$$

$$= \sum_{x=0}^{\infty} \frac{(x\lambda)^y e^{-x\lambda}}{y!} \frac{\Gamma(x + \phi^{-1})}{\Gamma(\phi^{-1})\Gamma(x+1)} \left(\frac{1}{1 + \mu\phi}\right)^{\phi^{-1}} \left(\frac{1}{1 + (\mu\phi)^{-1}}\right)^{x}$$

$$= f(y)\left(\frac{1}{1 + \mu\phi}\right)^{\phi^{-1}} + \sum_{x=1}^{\infty} \frac{(x\lambda)^y e^{-x\lambda}}{\Gamma(y+1)} \frac{\Gamma(x + \phi^{-1})}{\Gamma(\phi^{-1})\Gamma(x+1)} \left(\frac{1}{1 + \mu\phi}\right)^{\phi^{-1}} \left(\frac{1}{1 + (\mu\phi)^{-1}}\right)^{x}$$

where

$$f(y) = P(Y' = y | X = 0) = \begin{cases} 1, & y = 0 \\ 0, & y > 0 \end{cases}$$

**Estimating sensitivity**

Once the Maximum Likelihood Estimators (MLEs) are obtained for the log-likelihood function, it is possible to calculate the probability of $k$ fragments in a bin given observed $y$ read pairs according to Bayes' theorem:

$$P(X = k|Y' = y) = \frac{P(Y' = y|X = k)P(X = k)}{P(Y' = y)}$$

Then the probability that $n$ fragments carry the mutation among $k$ total fragments in the bin can be written as

$$\binom{k}{n} f^n(1 - f)^{k-n} = \frac{k!}{n!\,(k - n)!} f^n(1 - f)^{k-n}$$

Thus, the mutation frequency in the bin is $n/k$. An alternative way is to assume only one fragment has the mutation in the bin, considering the low frequency of the mutation.

Let $s$ denote the number of reads supporting the mutation. Given $y$ reads and the threshold that at least 3 reads support the mutation, the probability that the mutation cannot be detected is

$$P(s < 3;\ y) = \sum_{s=0}^{2} \binom{y}{s} (n/k)^s(1 - n/k)^{y-s}$$

If considering the sequencing error as $e$, it is possible to add the probability that 3 PCR products of the fragment support the mutation but at least one of them has a sequencing error, wherein the mutation cannot be detected:

$$P(s = 3;\ y) * (1 - (1 - e)^3) = \binom{y}{3} (n/k)^3(1 - n/k)^{y-3}(1 - (1 - e)^3)$$

Let $y_w$ and $y_c$ denote the number of read pairs in Watson and Crick families respectively. The probability that the mutation cannot be detected is

$$P(s < 3; y_w) + P(s < 3; y_c) - P(s < 3;\ y_w) * P(s < 3; y_c)$$

Then we can estimate the sensitivity at the position as

$$1 - \prod_{i=1}^{N_b} P(not\ detected\ in\ the\ i^{th}\ bin)$$

**Figure S1 Comparison between theoretical and empirical P-values from Monte Carlo procedures under truly distributed sequencing error rates**. With the null hypothesis, one million simulations were conducted.

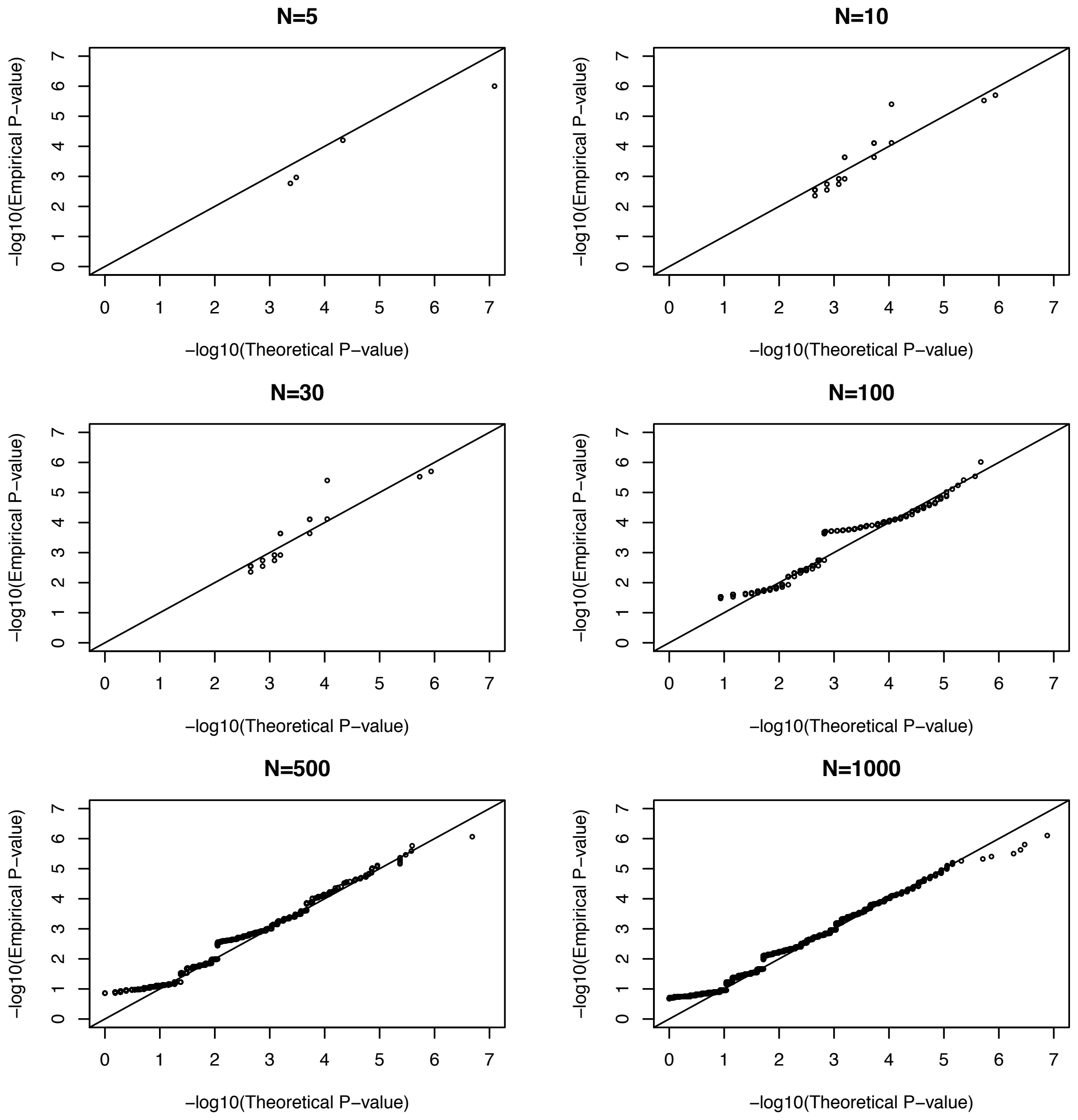

**Figure S2 Receiver Operating Characteristic (ROC) curves of the likelihood-based model with uniformly distributed sequencing errors (Q20).** We simulated 10,000 SNVs with a frequency equal to 1% or 0.1% and 1000 SNVs for 0.01%.

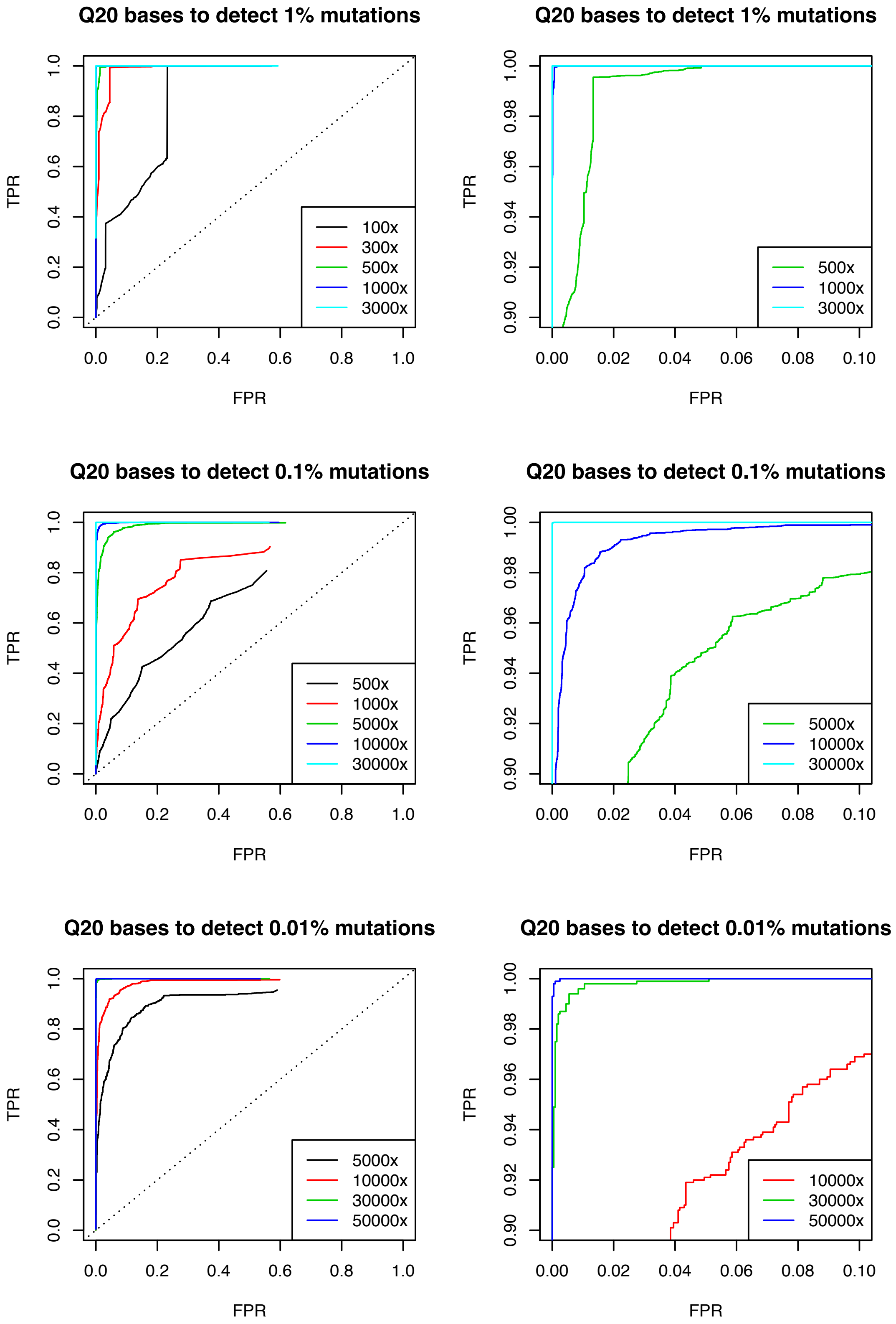

**Figure S3 Estimation of allele fraction with the likelihood-based model under uniformly distributed sequencing errors.** The differences between estimated and true fractions are consistent and predictable due to the nature of the likelihood-based model. The true fraction is always smaller than the estimated one. However, in practice, the accurate allele fraction is not as important as the existence of the allele.

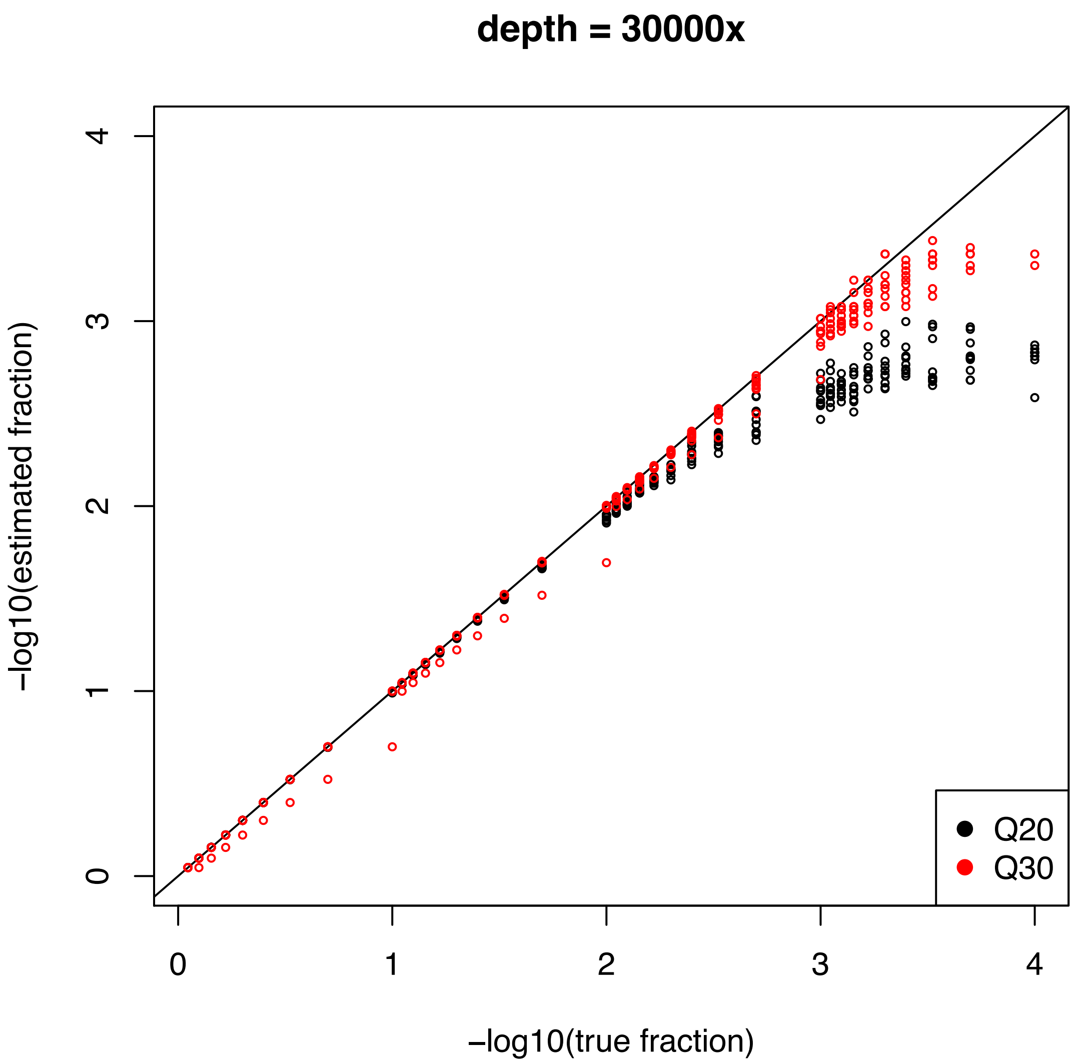

**Figure S4 Estimation of allele fraction with the likelihood-based model under true sequencing errors.** The results show that the allele fraction can be estimated accurately by using this model.

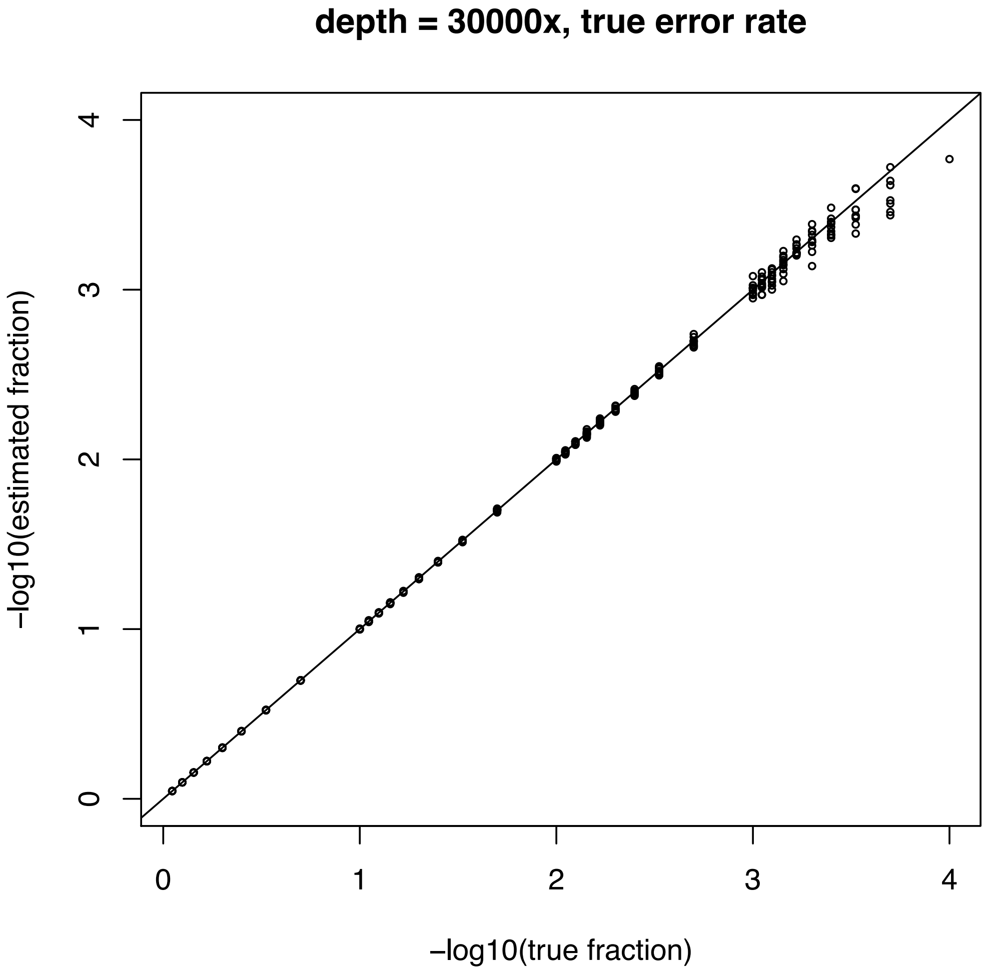

**Figure S5 Mutation spectrum of simulated data under different read depths (33k, 65k, 130k) without sensitivity adjustment.** It is obvious that the higher the read depth, the larger the number of detected mutations. The figure is based on all mutations without considering their mutation frequencies. In this case, as the original mutation lists are the same, the total numbers of mutations should be the same. This result indicates that sensitivity adjustment is essential when comparing multiple samples with different read depths.

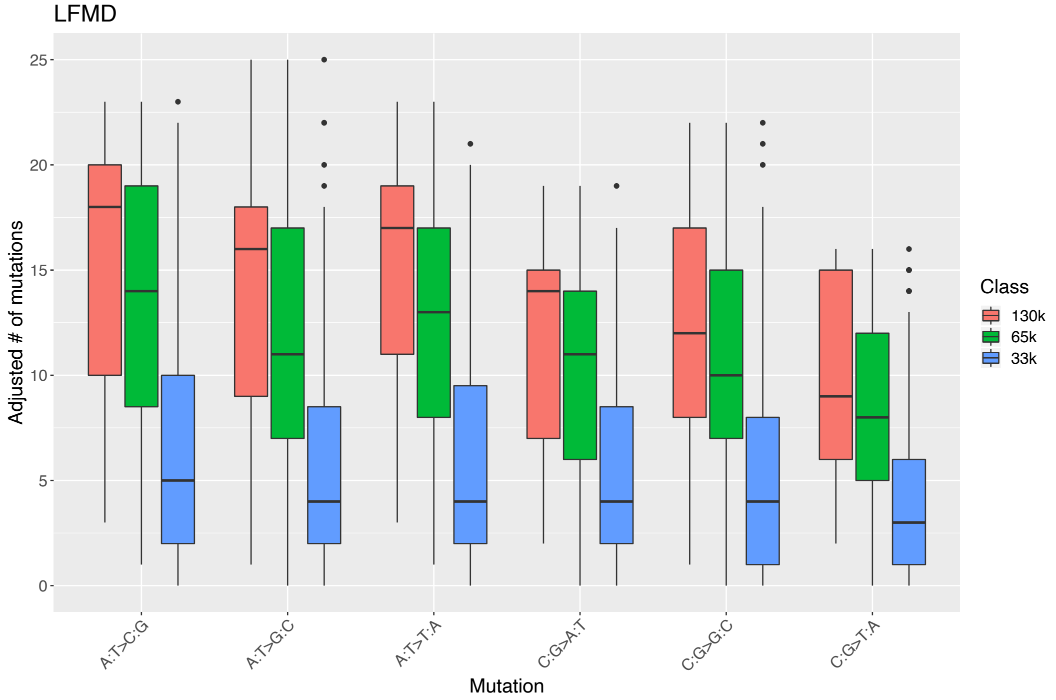

**Figure S6 Mutation spectrum of simulated data under different read depths (33k, 65k, 130k) with sensitivity adjustment.** The differences between 130k and 65k are smaller than in Figure S5, indicating that the sensitivity adjustment works well on samples with sufficient read depth.

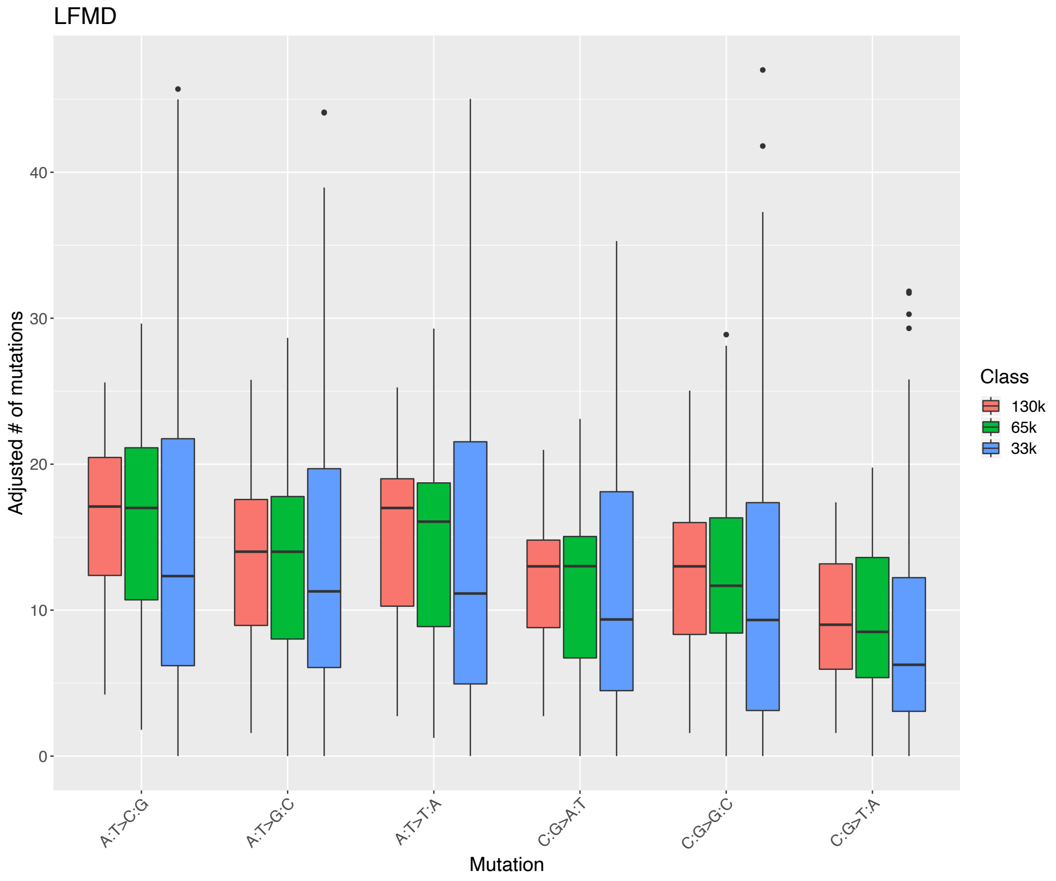

**Figure S7 Comparison of numbers of C:G>A:T mutations detected by four tools.** DS, Duplex Sequencing; UMI, UMI-tools; UniC, Unified Consensus Maker; LFMD, Low Frequency Mutation Detector; **, p-value < 0.01.

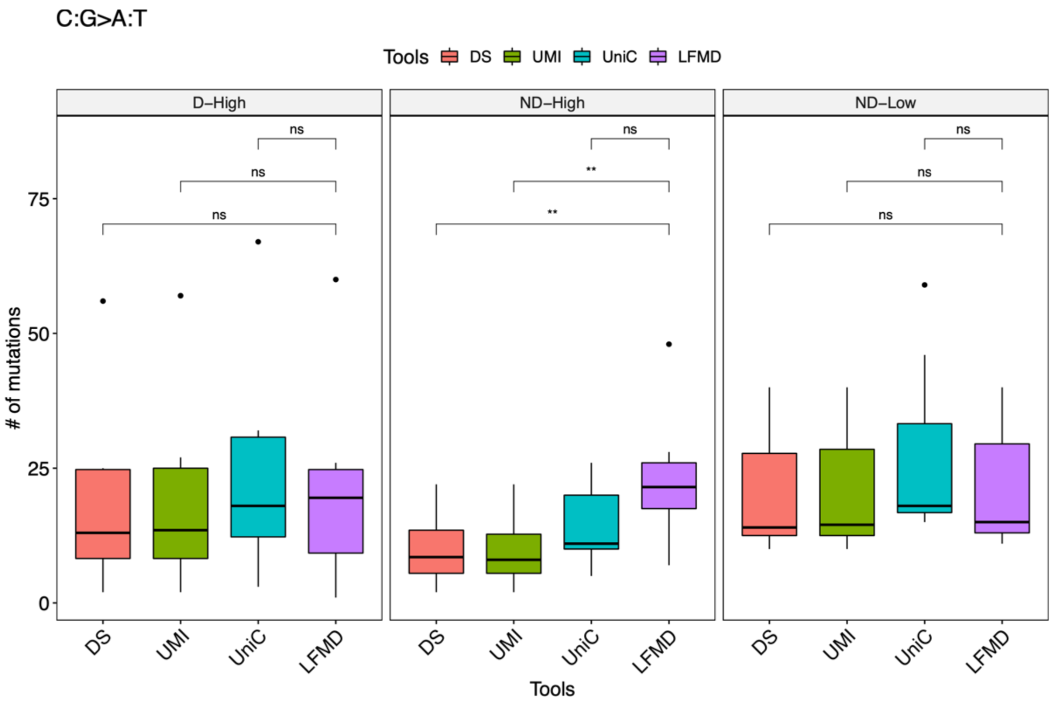


### References

1. Chen Y, Huang J, Ning Y et al. A conditional composite likelihood ratio test with boundary constraints, Biometrika 2017;105:225-232.
